# Transcriptome-wide association analysis of 211 neuroimaging traits identifies new genes for brain structures and yields insights into the gene-level pleiotropy with other complex traits

**DOI:** 10.1101/842872

**Authors:** Bingxin Zhao, Yue Shan, Yue Yang, Tengfei Li, Tianyou Luo, Ziliang Zhu, Yun Li, Hongtu Zhu

**Affiliations:** Department of Biostatistics, University of North Carolina at Chapel Hill, Chapel Hill, NC, USA; Department of Radiology, University of North Carolina at Chapel Hill, Chapel Hill, NC, USA; Biomedical Research Imaging Center, School of Medicine, University of North Carolina at Chapel Hill, Chapel Hill, NC, USA; Department of Genetics, University of North Carolina at Chapel Hill, Chapel Hill, NC, USA; Department of Computer Science, University of North Carolina at Chapel Hill, Chapel Hill, NC, USA

**Keywords:** Gene expression, Cross-tissue TWAS, Regional brain volumes, Diffusion tensor imaging, UK Biobank

## Abstract

Structural and microstructural variations of human brain are heritable and highly polygenic traits, with hundreds of associated genes founded in recent genome-wide association studies (GWAS). Using gene expression data, transcriptome-wide association studies (TWAS) can prioritize these GWAS findings and also identify novel gene-trait associations. Here we performed TWAS analysis of 211 structural neuroimaging phenotypes in a discovery-validation analysis of six datasets. Using a cross-tissue approach, TWAS discovered 204 associated genes (86 new) exceeding Bonferroni significance threshold of 1.37*10^−8^ (adjusted for testing multiple phenotypes) in the UK Biobank (UKB) cohort, and validated 18 TWAS or previous GWAS-detected genes. The TWAS-significant genes of brain structures had been linked to a wide range of complex traits in different domains. Additional TWAS analysis of 11 cognitive and mental health traits detected 69 overlapping significant genes with brain structures, further characterizing the genetic overlaps among these brain-related traits. Through TWAS gene-based polygenic risk scores (PRS) prediction, we found that TWAS PRS gained substantial power in association analysis compared to conventional variant-based PRS, and up to 6.97% of phenotypic variance (p-value=7.56*10^−31^) in testing datasets can be explained by UKB TWAS-derived PRS. In conclusion, our study illustrates that TWAS can be a powerful supplement to traditional GWAS in imaging genetics studies for gene discovery-validation, genetic co-architecture analysis, and polygenic risk prediction.

---

Brain structural and microstructural differences are phenotypically associated with many other complex traits across different categories, such as cognitive measures^1-5^, neurodegenerative/neuropsychiatric traits^6-9^, alcohol and tobacco consumption^10^, and physical bone density^11^. Structural variations of human brain can be quantified by multimodal magnetic resonance imaging (MRI). Specifically, the T1-weighted MRI (T1-MRI) can provide basic morphometric information of brain tissues, such as volume, surface area, sulcal depth, and cortical thickness. In region of interest (ROI)-based T1-MRI analysis, images are annotated onto ROIs of pre-defined brain atlas, and then both global (e.g., whole brain, gray matter, white matter) and local (e.g., basal ganglia structures, limbic and diencephalic regions) markers can be generated to measure the brain anatomy. On the other hand, diffusion MRI (dMRI) can capture local tissue microstructure through the random movement of water. Using diffusion tensor imaging (DTI) models, brain structural connectivity can be quantified by using white matter tracts extracted from dMRI, which build psychical connections among brain ROIs and are involved in connected networks for various brain functions^12,13^. See Miller, et al. ^11^ and Elliott, et al. ^14^ for a global overview and more information about neuroimaging modalities used in the present study.

Structural neuroimaging traits have shown moderate to high degree of heritability in both twin and population-based studies^14-24^. In the past ten years, genome-wide association studies (GWAS)^3,14,24-33^ have been conducted to identify the associated genetic variants (typically single-nucleotide polymorphisms [SNPs]) for brain structures. A highly polygenic^34,35^ genetic architecture has been observed, indicating that a large number of genetic variants contribute to the brain structure variations measured by neuroimaging biomarkers^21,36^. Particularly, using data from the UK Biobank (UKB^39^) cohort, two recent large-scale GWAS have identified 578 associated genes for 101 regional brain volumes derived from T1-MRI^37^ (referred as ROI volumes, n=19,629) and 110 DTI parameters of dMRI^38^ (referred as DTI parameters, n=17,706). Some of these discovered genes had been implicated with the same or other traits such as cognition and mental health diseases/disorders in previous GWAS. However, most of them have not been verified and need further investigations. As a supplement to traditional GWAS, recent advances of gene expression imputation methods^40-46^ and developments of reference databases (e.g., the Genotype-Tissue Expression (GTEx) project^47^) have put the transcriptome-wide association studies (TWAS) forward for gene-trait association analysis. Despite some challenges^48^ such as interpreting causality, TWAS has successfully discovered novel gene-trait associations and provided new insights into biological mechanisms for many complex traits^49^. Through imputed transcriptomes, TWAS can reduce the multiple testing burden and leverage gene expression data to increase testing power for gene-trait association detection. This is a particularly desirable feature for imaging genetics studies, for which most of neuroimaging GWAS datasets continue to have small sample sizes and heavy multiple testing burden^50^.

Here we applied TWAS methods to 211 structural neuroimaging traits including 101 ROI volumes and 110 DTI parameters. As these brain-related traits tend to be highly polygenic^21,36^ and are related with many traits across different categories^11^, we used a cross-tissue (panel) TWAS approach (UTMOST^42^) in our main analysis. UTMOST first performs single-tissue gene-trait association analysis in each reference panel with both within-tissue and cross-tissue statistical penalties, and then combines these single-tissue results using the Generalized Berk-Jones (GBJ) test^51^, which is aware of tissue-dependence and can account for the potential sharing of local expression regulation across tissues. The UKB dataset was used in the discovery phase (n=19,629 for ROI volumes and 17,706 for DTI parameters, respectively). For the same UKB cohort, we compared TWAS-significant genes to previous GWAS findings in gene-based association analysis via MAGMA^52^ and gene-level functional mapping and annotation results by FUMA^53^. The UKB TWAS results were validated in five independent data sources, including Philadelphia Neurodevelopmental Cohort (PNC^54^,n=537), Alzheimer’s Disease Neuroimaging Initiative (ADNI^55^, n=860), Pediatric Imaging, Neurocognition, and Genetics (PING^56^, n=461), the Human Connectome Project (HCP^57^, n=334), and the ENIGMA2^24^ and ENIGMA-CHARGE collaboration^33^ (n=13,193, for 8 ROI volume traits, referred as ENIGMA in this paper). Additional TWAS analysis was performed on 11 cognitive and mental traits to explore their genetics overlaps with brain structures. Chromatin interaction enrichment analysis and drug-target lookups were conducted for TWAS-significant genes. Finally, we developed TWAS gene-based polygenic risk scores^58^ (PRS) using FUSION^40^ to fully assess polygenic architecture and examine the predictive ability of the UKB TWAS results.

## RESULTS

### Overview of TWAS discovery-validation in the six datasets

We conducted a two-phase discovery-validation TWAS analysis for 211 neuroimaging traits by using the UKB cohort for discovery and the other datasets (ADNI, HCP, PING, PNC, and ENIGMA) for validation. We applied the UTMOST gene expression imputation models trained on 44 GETx (v6) reference panels, and used GWAS summary statistics generated from previous GWAS as inputs. In the rest of this paper, we refer 1.37*10^−8^ (that is, 5*10^−2^/17,290/211, adjusted for all candidate genes and traits performed) as the significance threshold for gene-trait associations unless otherwise stated.

The UKB discovery phase identified 614 significant gene-trait associations (**Supplementary Table 1**) between 204 genes and 135 neuroimaging traits (53 ROI volumes, 82 DTI parameters). Of the 204 TWAS-significant genes, 61 (29.9%) had significant associations with more than two neuroimaging traits, 25 (12.3%) had more than five significant associations, and 12 (5.9%) had at least ten, including *OSER1, XRCC4, PLEKHM1, ZKSCAN4, EIF4EBP3, MAPT, LRRC37A, CRHR1, FOXF1, TREH, ARHGAP27*, and *C6orf100*. These 12 genes together contributed 195 (31.8%) of the 614 gene-trait associations, indicating their widespread influences on brain structures. Specifically, we identified 123 genes whose imputed gene expression levels were significantly associated with one of more of the 53 ROI volumes (215 associations in total, 115 new, **Supplementary Fig. 1**), and 103 significantly associated genes (22 overlapping) for one or more of the 82 DTI parameters (399 associations in total, 219 new, **Supplementary Fig. 2**). **Figure 1** illustrates that TWAS prioritized previous GWAS findings of MAGMA and FUMA and also discovered many new associations and genes. Moreover, some genes were associated with both ROI volumes and DTI parameters, while others were more specifically related to certain structures (**Supplementary Fig. 3**). For example, *XRCC4, ZKSCAN4, EIF4EBP3*, and *CD14* were associated with DTI parameters but not ROI volumes, *DEFB124, COX4I2, HCK, HM13*, and *REM1* showed associations with putamen and pallidum volumes, and the associations of *PLEKHM1, LRRC37A, MAPT, CNNM2, NT5C2, ARHGAP27*, and *CRHR1* were spread widely across DTI parameters and total brain volume.

**Figure 1.**
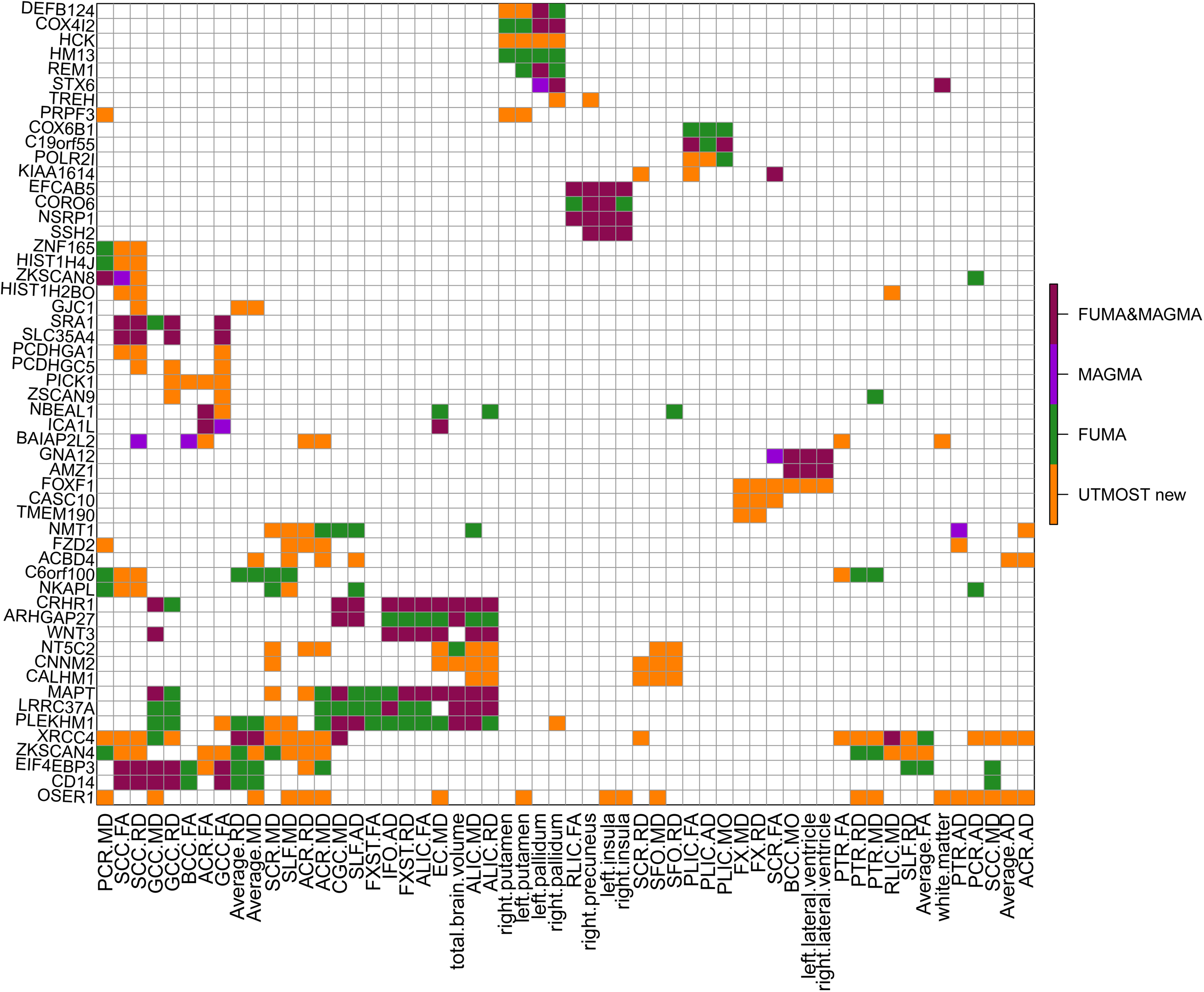
Selected significant gene-trait associations discovered in UKB (UK Biobank) cross-tissue TWAS analysis of 211 neuroimaging traits (n=19,629 subjects for ROI volumes and 17,706 for DTI parameters). The gene-level associations were estimated and tested by the cross-tissue UTMOST approach (https://github.com/Joker-Jerome/UTMOST). We used the p-value threshold of 1.37*10^−8^, corresponding to adjusting for testing 211 imaging phenotypes with the Bonferroni correction. The *x* axis provides the IDs of the neuroimaging traits, and the *y* axis lists the detected genes in TWAS. The new (UTMOST new) and previously reported GWAS-significant associations (MAGMA, FUMA, and FUMA&MAGMA) were labeled with different colors (orange, purple, green, and red, respectively).

We validated the UKB results in the other five independent cohorts. For each dataset, we applied the Bonferroni-corrected significance threshold accounting for all candidate genes and traits analyzed (that is, 5*10^−2^/17,290/number of traits, **Supplementary Tables 2-6**). We found that 13 UKB TWAS-significant genes and 5 more previous GWAS-significant genes can be validated in one or more of the five validation datasets (**Supplementary Fig. 4**) including *ANKRD42, DCC, DCTPP1, DLGAP5, HCK, LGALS3, UBE2C, KLRD1, LRRC37A, OSER1, PRPF3, TREH, TGM7, NUP210L, DOK5, KRTAP5-1, C20orf166*, and *DPP4*. The TWAS novel findings and validated genes were discussed further in details below.

### Novel TWAS discoveries and validated genes

Of the 204 UKB TWAS-significant genes, 90 were not discovered in previous GWAS of the same UKB dataset (**Supplementary Table 7**). TWAS resulted in 60 new associated genes for 53 ROI volumes (106 associations, **Supplementary Fig. 5**), and 52 new genes for 82 DTI parameters (139 associations, **Supplementary Fig. 6**). According to NHGRI-EBI GWAS catalog^59^, the 90 TWAS-significant genes replicated four previous findings on brain structures, including *JPH3*^60^ for hippocampal volume in mild cognitive impairment, *CNNM2*^61^ for white matter lesion progression, *FOXF1*^62^ for hippocampal volume in Alzheimer’s disease progression, and *C1QL1*^63^ for white matter hyperintensity burden. The other 86 genes had not been linked to brain structure previously and thus can be regarded as novel genes for these 211 neuroimaging traits. To explore the genetic overlaps with other traits in different domains, we performed association lookups for the 90 TWAS-significant genes on the NHGRI-EBI GWAS catalog (**Supplementary Table 8**)**. Figure 2** shows that these genes were widely associated with physical measures (e.g., height, waist-to-hip ratio, heel bone mineral density, body mass index), cognitive traits (e.g., cognitive function, intelligence, math ability), neuropsychiatric and neurodegenerative diseases/disorders (e.g., schizophrenia, bipolar disorder, Alzheimer’s disease), coronary artery disease, mean corpuscular hemoglobin, neuroticism, education, reaction time, chronotype, smoking behavior and alcohol use, such as *CDK2AP1*^*64-67*^, *ELL*^*68-70*^, *CTTNBP2*^*71-73*^, and *SH2B1*^*72,74-76*^.

**Figure 2.**
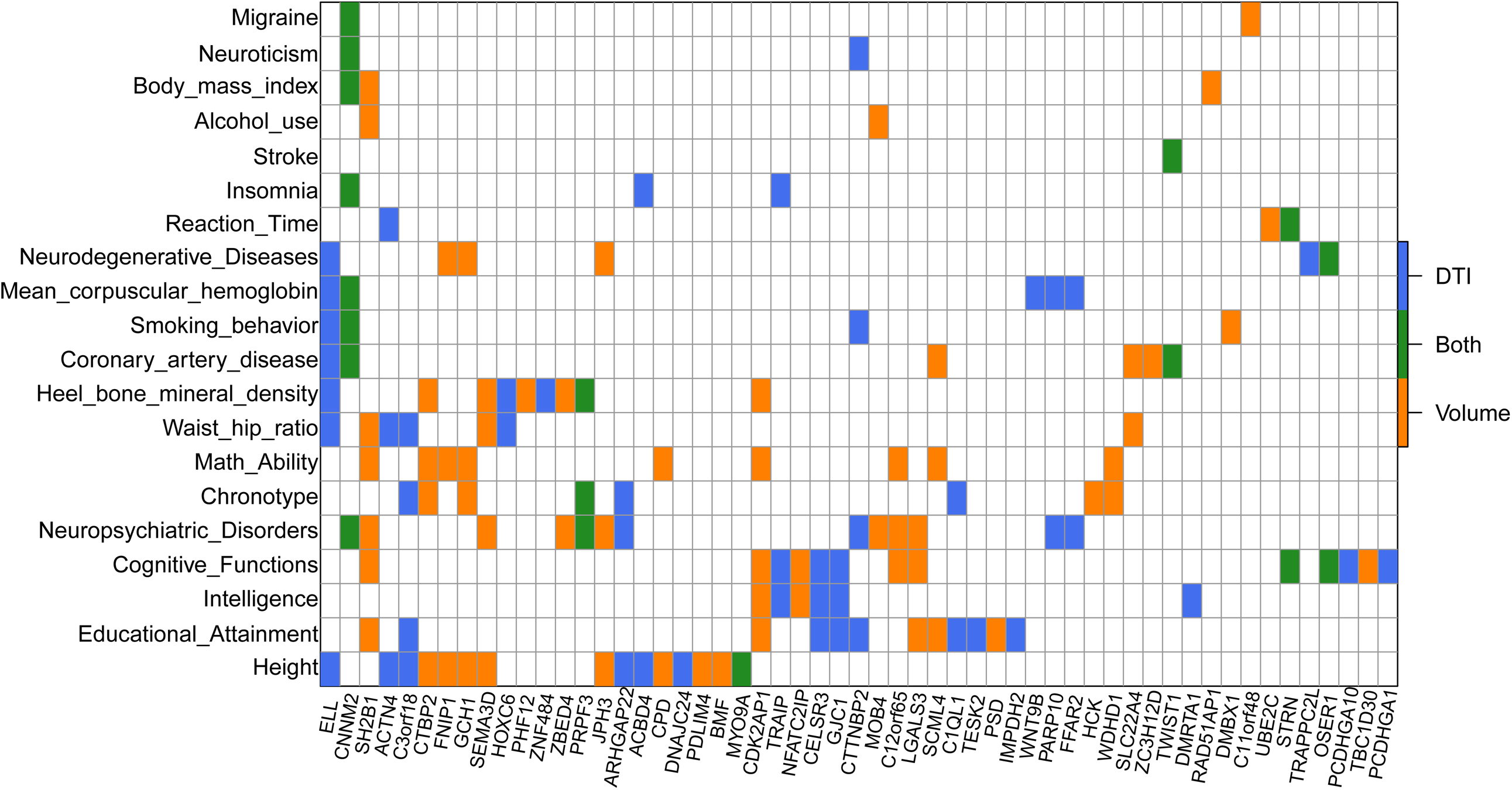
TWAS-significant genes of neuroimaging traits (n=19,629 subjects for ROI volumes and 17,706 for DTI parameters) that have been linked to other complex traits in previous GWAS. For each of the TWAS-significant genes listed in the *x* axis, we manually checked the previously reported associations on the NHGRI-EBI GWAS catalog (https://www.ebi.ac.uk/gwas/). The genes associated with DTI parameters (DTI), ROI volumes (Volume), and both of them (Both) were labeled with three different colors (blue, orange, and green, respectively).

For the 18 TWAS-validated genes shown in **Supplementary Fig. 4**, 8 (*ANKRD42, DCC, LRRC37A, NUP210L, DOK5, KRTAP5-1, C20orf166*, and *DPP4*) of them had been discovered in the previous UKB GWAS and were implicated in brain-related complex traits, such as neuroticism^64^, major depression^77^, schizophrenia^78-80^, Intelligence^81^, math ability^73^, reaction time^75^, and insomnia^82^. The left ten genes, which were novel findings of TWAS, also had known associations with many cognitive and mental health traits. For example, previous GWAS reported that *HCK* was associated with chronotype^82^, *LGALS3* with schizophrenia^83^, *UBE2C* with reaction time^75^, *KLRD1* with adolescent idiopathic scoliosis^84^, *OSER1* with cognitive performance^77^ and Alzheimer’s disease^76^, and *PRPF3* with chronotype^76,85^ and neuropsychiatric disorders^86^. In summary, TWAS novel and validated genes expand the overview of gene-level pleiotropy across these traits, suggesting that neuroimaging-derived biomarkers could be useful in studying a wide range of complex traits.

### Compared to brain tissue-specific TWAS analysis

As a comparison, we performed a brain tissue-specific version of TWAS that only combines brain tissues in UTMOST (Method). This brain tissue-specific TWAS detected 308 significant gene-trait associations (**Supplementary Table 9**) between 107 unique genes and 96 neuroimaging traits, including 64 associated genes for one or more of 37 ROI volumes (104 associations, **Supplementary Fig. 7**), and 53 genes (10 overlapping) for one or more of 59 DTI parameters (204 associations, **Supplementary Fig. 8**).

Most (101/107) of the tissue-specific genes have been identified by either the cross-tissue TWAS (95/107) or previous GWAS (70/107). The 6 genes that were uniquely identified by tissue-specific analysis included *KNCN, LHFPL3, MBD2, TBK1, C3orf62*, and *TMEM173. LHFPL3* showed associations with education^87^, social behavior^88,89^, cognitive ability^75^, schizophrenia^90^, and bipolar disorder^91^. *MBD2* was associated with reaction time^75^, *TBK1* with amyotrophic lateral sclerosis^92,93^, and *C3orf62* with intelligence^82^. Compared to tissue-specific TWAS, cross-tissue analysis clearly identified more signals.

For example, of the 215 gene-trait associations identified by cross-tissue analysis of ROI volumes, 100 had been identified in GWAS, 28 can be additionally identified by tissue-specific TWAS, and 87 can only be detected by cross-tissue analysis (**Supplementary Fig. 9**). Similarly, 180 of the 399 cross-tissue TWAS associations for DTI can be identified in GWAS, 69 can be additionally identified by tissue-specific TWAS, and 150 were cross-tissue TWAS only (**Supplementary Fig. 10**). These results illustrate the advantage of cross-tissue analysis over brain tissue-specific TWAS for discovering association signals that are difficult to be identified in traditional GWAS. We further compared their results in a few follow-up analyses below.

### Comparison with GWAS variant-level signals and conditional analysis

For each of the 614 gene-trait associations detected in cross-trait TWAS, we used previous GWAS summary statistics to check the most significant variant within the gene region (with a 1MB window on each side) that was pinpointed in the same UKB dataset (Method). The GWAS p-value of the most significant variant was greater than 1*10^−6^ for any associations of 13 genes (**Supplementary Table 10**). None of them had been identified by MAGMA or FUMA, indicating that it can be difficult to detect these genes by GWAS or post-GWAS screening for any of these neuroimaging traits. Of the 13 genes, 7 (*OSER1, TREH, PRPF3, KLRD1, TGM7, DCTPP1, UBE2C*) were validated in one or more of the five validation datasets and were discussed in previous section. For the other 6 genes (*CELSR3, MYO9A, DNAJC24, GYPE, TMEM136, MOB4*) genes, *MOB4* was reported for major depression^94^ and autism spectrum disorder/schizophrenia^95^, *DNAJC24* was linked to adolescent idiopathic scoliosis^84^, and *CELSR3* was associated with education^65^ and cognitive ability^64,81^. The same checking was then performed for the 308 significant gene-trait associations of brain tissue-specific TWAS. We found that only one gene *DCTPP1* had minimum GWAS p-value greater than 1*10^−6^ (**Supplementary Table 11**).

We next performed a conditional analysis to see whether the TWAS signals remained significant after adjustment for the most significant genetic variant used in UTMOST gene expression imputation models (Method). Although our cross-tissue analysis combined information from many genetic variants across various human tissues, we found that 418 of the 614 associations may indeed be dominated by the strongest GWAS signal of the imputation model, as their conditional p-values were larger than 0.05 (**Supplementary Table 12**). However, the conditional p-values of four genes (*XRCC4, OBFC1, C15orf56, NMT1*) were smaller than 1*10^−6^ for 18 gene-trait associations, suggesting that these associations were unlikely to be driven by a signal genetic variant. When the p-value threshold was relaxed to 1*10^−3^, 66 associations of 20 genes persisted after conditional analysis. The conditional analysis was also performed on significant associations of brain tissue-specific TWAS. Their conditional p-values were smaller than 1*10^−6^ for three genes (*XRCC4, C15orf56, NMT1*) with 15 associations, and were smaller than 1*10^−3^ for 10 genes with 42 associations (**Supplementary Table 13**).

### Additional TWAS analysis for cognitive and mental health traits

To further explore the gene-level genetic overlaps among brain structure and other brain-related traits, we performed cross-tissue TWAS analysis for 11 cognitive and mental health traits (**Supplementary Table 14**). We found that 69 of the 204 TWAS-significant genes of neuroimaging traits were also significantly associated with one or more of the 11 cognitive and mental health traits (**Figure 3**). These results suggest the genes involved in brain structure changes are often also active in brain functions and mental disorder/diseases. For example, we found 33 overlapping genes with cognitive function, 32 with education, 26 with numerical reasoning, 25 with intelligence, 23 with neuroticism, 19 with drinking behavior, and 13 with schizophrenia. A large proportion (48/69) of these genes were associated with more than one cognitive or mental health traits, and 11 genes were linked to at least five traits, including *SCML4, C16orf54, DCC, NFATC2IP, NPIPB7, NPIPB9, SH2B1, CRHR1, LRRC37A, HIST1H2BO*, and *NKAPL*, indicating the high degree of statistical pleiotropy^96^ of these genes.

**Figure 3.**
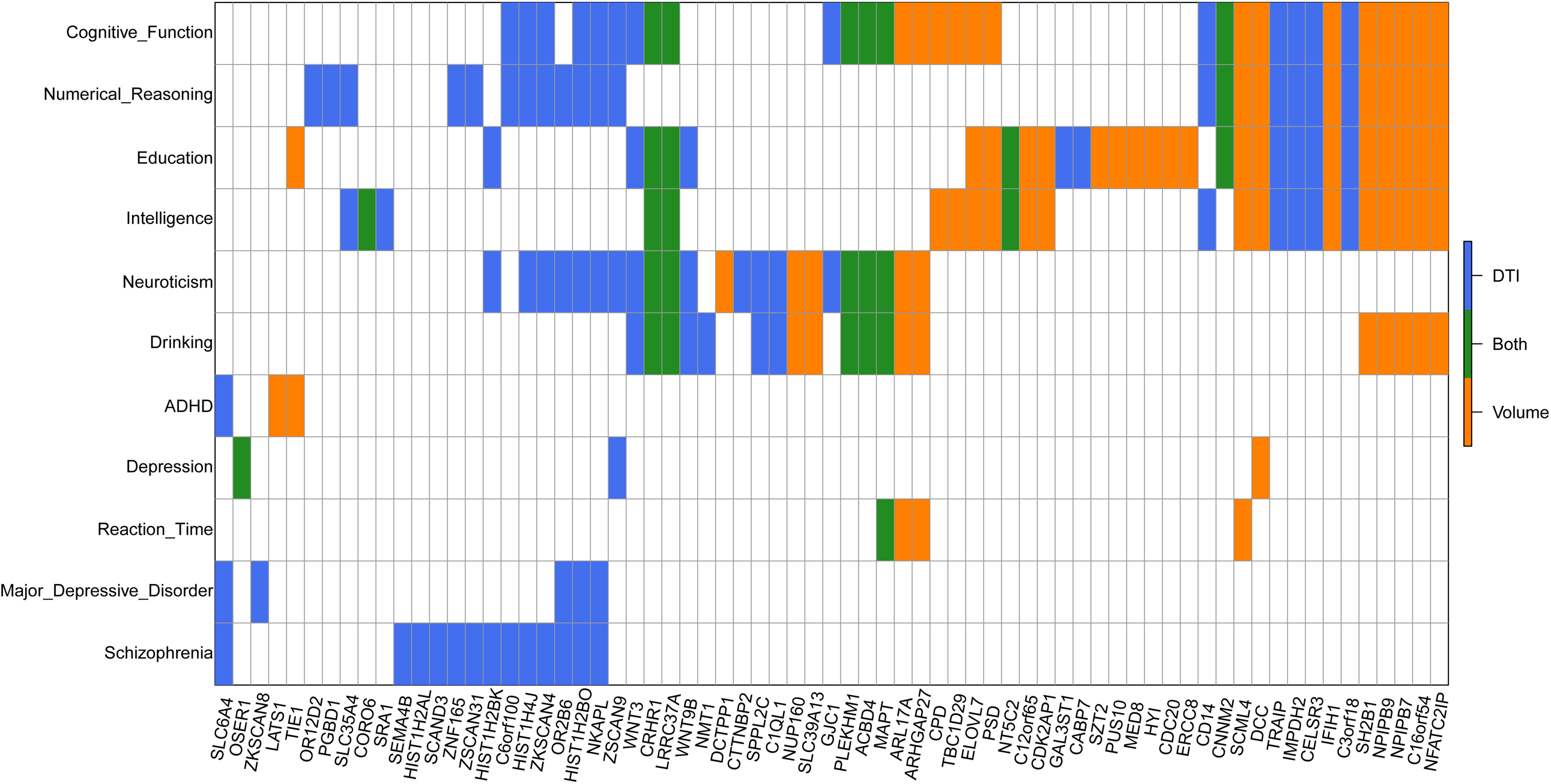
Overlapping TWAS-significant genes between neuroimaging traits (n=19,629 subjects for ROI volumes and 17,706 for DTI parameters) and 11 cognitive and mental health traits. The gene-level associations were estimated and tested by the cross-tissue UTMOST approach (https://github.com/Joker-Jerome/UTMOST). We adjusted for testing 211 neuroimaging traits (p-value threshold 1.37*10^−8^) and 11 cognitive traits (p-value threshold 2.63*10^−7^) with the Bonferroni correction, respectively. The *x* axis provides the IDs of the neuroimaging traits. The *y* axis lists the 11 cognitive and mental health traits, and **Supplementary Table 23** details the resources of their GWAS summary statistics and the sample sizes of corresponding studies.

**Figure 4.**
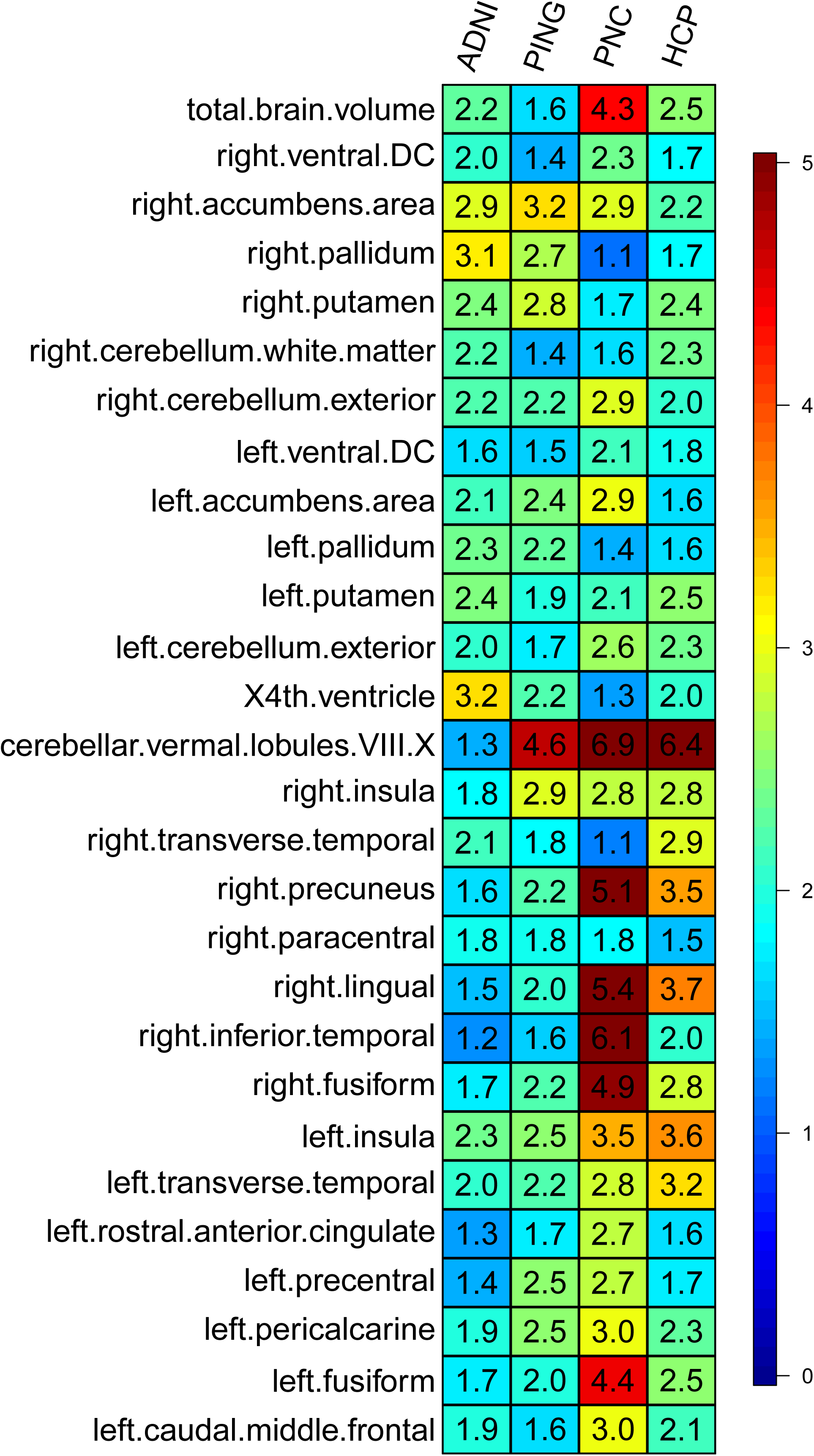
Prediction accuracy (incremental R-squared) of gene-based polygenic risk scores constructed by UKB TWAS results (n=19,629 subjects) on the four independent datasets. The *x* axis lists the four independent cohorts (ADNI, HCP, PING and PNC) and the *y* axis lists the ROI volumes. The displayed numbers are the proportions of phenotypic variation that can be additionally explained by UKB TWAS-derived gene-based PRS.

### Chromatin interaction enrichment analysis and drug-target lookups

To explore the biological interpretations of TWAS and GWAS-significant genes, we performed enrichment analysis in promoter-related chromatin interactions of four types of brain neurons^97^ (iPSC-induced excitatory neurons, iPSC-derived hippocampal DG-like neurons, iPSC-induced lower motor neurons, and primary astrocytes), and also in high confident interactions of adult and fetal cortex^98^ (Method). The raw p-values of Wilcoxon rank test for enrichment were summarized in **Supplementary Table 15.** We found that cross-tissue TWAS-significant genes of the 11 cognitive and mental health were significantly enriched in chromatin interactions from all of the five validation datasets (p-value range=[4.91*10^−11,^ 3.03*10^−5^]), suggesting that TWAS-significant genes actively interacted with other chromatin regions and played a more important role in regulating gene expressions as compared with other genes. The cross-tissue TWAS-significant genes of neuroimaging traits also showed significant enrichments (p-value range= [1.38*10^−3^,2.44*10^−2^]). Merging the two sets of genes resulted in smaller p-value in each dataset (p-value range=[2.93*10^−11,^ 2.77*10^−5^]). The most significant enrichment was observed in iPSC-induced lower motor neurons. These results remained significant after adjusting for multiple testing by using Benjamini-Hochberg (B-H) procedure at 0.05 level (**Supplementary Table 16**). In contrast, GWAS-significant genes were only significantly enriched in primary astrocytes and high confident interactions (p-value range=[5.11*10^−3,^ 1.48*10^−2^]), and brain tissue-specific TWAS-significant genes did not show any significant enrichments after B-H adjustment.

We carried out drug-target lookups using a recently published drug-target database^99^ to see whether any of the TWAS and GWAS-significant genes were known targets of existing drugs. We focused on nervous system drugs with Anatomical Therapeutic Chemical (ATC) code started with “N”, yielding 2,285 drug-gene pairs between 273 drugs and 241 targeted genes. We found that 12 TWAS-significant genes of the 11 cognitive and mental health traits were known targets for 64 drugs, including *CACNA1I, ESR1, ALDH2, CACNA1C, GRM2, KCNJ3, SCN3A, CACNA1D, KCNK3, CHRNA3, CHRNA6*, and *SLC6A4*. Of the 64 drugs, 27 were anti-depressants (ATC: N06A) to treat major depressive disorder and other conditions, and 10 were anti-psychotics (ATC: N05A) to manage psychosis such as schizophrenia and bipolar disorder (**Supplementary Table 17**). In addition, 3 more drug-target genes (*GABBR1, HTR2B, CREB1*) were detected by GWAS or TWAS of neuroimaging traits (**Supplementary Table 18**). These 3 genes were targets for 19 more drugs, 6 of which were anti-Parkinson drugs (ATC: N04) for treatment of Parkinson’s disease and related conditions, and 5 were anti-migraine preparations (ATC: N02C) used in prophylaxis and treatment of migraine. These results may suggest that TWAS-significant genes could be considered as new targets in future drug development.

### TWAS gene-based polygenic risk scores analysis

To fully assess the polygenic genetic architecture of neuroimaging traits and examine the predictive ability of UKB TWAS results, we constructed TWAS gene-based PRS on subjects in PNC, HCP, PING, and ADNI cohorts for all of the 211 neuroimaging traits (Method). The prediction analysis was conducted separately on 52 reference panels (13 GETx v7 brain tissues, 35 GTEx v7 other tissues, 1 non-GETx brain tissue, and 3 non-GETx other tissues) using the FUSION^40^ software and database. We found that genetically predicted profiles for 28 ROI volumes (**Figure 5**) and 23 DTI parameters (**Supplementary Fig. 11**) were significantly associated with the corresponding observed traits in all testing datasets after Bonferroni correction (that is, 101*4+3*110=734 tests). Compared to previous SNP-based PRS analysis that yielded significant PRS profiles for 11 ROI volumes^37^, gene-based PRS profiles were significant for more ROI volumes, such as left/right insula, left/right pallidum, left/right ventral DC, left/right fusiform, and left/right transverse temporal, suggesting the substantial power gain in association analysis of PRS. The significant TWAS PRS can account for 0.97%-6.97% phenotypic variance (p-value range=[8.0*10^−29^, 6.81*10^−5^]) (**Supplementary Tables 19-20**), which was within the similar range to SNP-based PRS analysis. For example, the (incremental) R-squared of TWAS PRS of Cerebellar vermal lobules VIII–X was 6.97% in PNC and 6.48% in HCP, and the R-squared of SFO MD-derived TWAS PRS was 3.8% in PING and 2.41% in PNC. We also examined the performance of each reference panel on these significant traits. There was a significant linear relationship between the panel sample size and average prediction R-squared (48 GTEx reference panels, simple correlation=0.53, p-value=1.21*10^−4^, **Supplementary Fig. 12**), which means that currently panel sample size may dominate the performance of TWAS PRS analysis regardless of the tissue specificity^58^. Among the brain tissue panels, we found that cerebellum tissue had the largest sample size and also showed the highest average R-squared (**Supplementary Table 21**), further supporting the importance of reference panel sample size.

## DISCUSSION

In this study, we applied TWAS methods on 211 neuroimaging traits to identify genes, whose imputed expression levels were associated with brain structure variations. Using a cross-tissue approach, our main discovery analysis identified 86 novel genes and validated 18 significant genes at stringent Bonferroni-correction p-value thresholds. Conditional analysis and comparison with GWAS variant-level results suggested that the identification and validation of new genes reflect the ability of TWAS to reduce the testing burden and to combine the small genetic variant effects. We also performed brain tissue-specific TWAS and illustrated the unique strengths of cross-tissue TWAS in conditional and enrichment analyses. Lots of brain structure-related genes were known genetic factors for a wide range of complex traits, ranging from physical traits, cognition, mental disease/disorders, blood assays, to lifestyle, which extend the potential applications of neuroimaging traits. Some of these genetic overlaps were additionally highlighted by a TWAS analysis of 11 cognitive and mental health traits.

The present study faces some limitations. First, since these results are purely based on statistical associations, it is hard to draw conclusions about the underlying causality and prioritize causal genes^42,100^. This is also one of the main challenges for most of the current TWAS approaches^48^. Follow-up experimental validation is a clear need to confirm TWAS results and pinpoint the causal genes of brain structure changes. Second, the brain tissue-specific TWAS did not yield much new results compared to the previous GWAS and brain tissue panels did not show better prediction accuracy than non-brain tissues in gene-based PRS analysis. Both of the two observations support the use of multiple tissues in our analysis to increase testing power for association analysis, but making the causality interpretation of TWAS results even more complicated. In addition, though gene-based PRS had much better power in association tests than SNP-based polygenic scores, their prediction accuracies were similar. These limitations may be due to the fact that currently brain tissue reference panels do not have large sample size and/or the associated gene expression imputations may have low quality. Despite these limitations, it is clear that TWAS have the potential to become a powerful supplement to traditional GWAS in imaging genetics studies. In our study, many new gene-trait associations were discovered and the underlying genetic overlaps among complex traits were largely expanded. With better brain tissue gene expression reference panels and more neuroimaging GWAS datasets available, future TWAS analyses of neuroimaging traits are expected to show the value of tissue specificity and improve our understanding for the genetic basis of human brain.

## ACKNOWLEDGEMENTS

This research was partially supported by U.S. NIH grants MH086633 (H.Z.) and MH116527 (TF.L.). We thank Quan Wang, Bingshan Li, and Jia Wen for helpful conversations. We thank the individuals represented in the UK Biobank, ADNI, HCP, PING, PNC, ENIGMA2, and ENIGMA-CHARGE datasets for their participation and the research teams for their work in collecting, processing and disseminating these datasets for analysis. This research has been conducted using the UK Biobank resource (application number 22783), subject to a data transfer agreement. We gratefully acknowledge all the studies and databases that made GWAS summary data available. Part of data collection and sharing for this project was funded by the Alzheimer’s Disease Neuroimaging initiative (ADNI) (National Institutes of Health Grant U01 AG024904) and DOD ADNI (Department of Defense award number W81XWH-12-2-0012). ADNI is funded by the National Institute on Aging, the National Institute of Biomedical Imaging and Bioengineering and through generous contributions from the following: Alzheimer’s Association; Alzheimer’s Drug Discovery Foundation; Araclon Biotech; BioClinica, Inc.; Biogen Idec Inc.; Bristol-Myers Squibb Company; Eisai Inc.; Elan Pharmaceuticals, Inc.; Eli Lilly and Company; EuroImmun; F. Hoffmann-La Roche Ltd and its affiliated company Genentech, Inc.; Fujirebio; GE Healthcare; IXICO Ltd; Janssen Alzheimer Immunotherapy Research & Development, LLC; Johnson & Johnson Pharmaceutical Research & Development LLC; Medpace, Inc.; Merck & Co., Inc.; Meso Scale Diagnostics, LLC; NeuroRx Research; Neurotrack Technologies; Novartis Pharmaceuticals Corporation; Pfizer Inc.; Piramal Imaging; Servier; Synarc Inc.; and Takeda Pharmaceutical Company. The Canadian Institutes of Health Research is providing funds to support ADNI clinical sites in Canada. Private sector contributions are facilitated by the Foundation for the National Institutes of Health (www.fnih.org). The grantee organization is the Northern California Institute for Research and Education, and the study is coordinated by the Alzheimer’s Disease Cooperative Study at the University of California, San Diego. ADNI data are disseminated by the Laboratory for Neuro Imaging at the University of Southern California. Part of the data collection and sharing for this project was funded by the Pediatric Imaging, Neurocognition and Genetics Study (PING) (U.S. National Institutes of Health Grant RC2DA029475). PING is funded by the National Institute on Drug Abuse and the Eunice Kennedy Shriver National Institute of Child Health & Human Development. PING data are disseminated by the PING Coordinating Center at the Center for Human Development, University of California, San Diego. Support for the collection of the PNC datasets was provided by grant RC2MH089983 awarded to Raquel Gur and RC2MH089924 awarded to Hakon Hakonarson. All PNC subjects were recruited through the Center for Applied Genomics at The Children’s Hospital in Philadelphia. HCP data were provided by the Human Connectome Project, WU-Minn Consortium (Principal Investigators: David Van Essen and Kamil Ugurbil; 1U54MH091657) funded by the 16 NIH Institutes and Centers that support the NIH Blueprint for Neuroscience Research; and by the McDonnell Center for Systems Neuroscience at Washington University.

## AUTHOR CONTRIBUTIONS

B.Z., Y.S., Y.L., and H.Z. designed the study. B.Z., Y.S., Y.Y., TF.L., TY.L., and Z.Z performed the experiments and analyzed the data. B.Z., Y.S., Y.L., and H.Z. wrote the manuscript with feedback from all authors.

## COMPETETING FINANCIAL INTERESTS

The authors declare no competing financial interests.

## METHODS

### GWAS summary statistics datasets

We made use of GWAS summary statistics to test for gene-trait associations in our TWAS study. The GWAS summary-level were from six studies, including the UK Biobank (UKB, http://www.ukbiobank.ac.uk/resources/) study, the Human Connectome Project (HCP, https://www.humanconnectome.org/) study, the Pediatric Imaging, Neurocognition, and Genetics (PING, http://www.chd.ucsd.edu/research/ping-study.html) study, the Philadelphia Neurodevelopmental Cohort (PNC, https://www.ncbi.nlm.nih.gov/projects/gap/cgi-bin/study.cgi?study_id=phs000607.v1.p1) study, the Alzheimer’s Disease Neuroimaging Initiative (ADNI, http://adni.loni.usc.edu/data-samples/) study, and ENIGMA2 (GWAS of subcortical volumes) and the ENIGMA-CHARGE collaboration (http://enigma.ini.usc.edu/research/). For discovery, we used the GWAS summary statistics of the UKB study. Then the GWAS results of the other studies were used for validation, see **Supplementary Table 22** for a summary of sample size and the analyzed neuroimaging traits of each GWAS. More information about study cohorts and neuroimaging traits can be found in the original GWAS^24,33,37,38^. We also performed TWAS analysis for 11 cognitive and mental health traits, see **Supplementary Table 23** for these data resources.

### Cross-tissue TWAS analysis by UTMOST

Cross-tissue TWAS analysis was performed for each trait using the UTMOST software (https://github.com/Joker-Jerome/UTMOST). We first run single-tissue association test for each of the 44 GTEx (v6) reference panels using the above GWAS summary statistics as input. There were 17,290 candidate genes considered in UTMOST. Second, the gene-trait associations in 44 panels (tissues) were combined by the GBJ test (https://cran.r-project.org/web/packages/GBJ/). We used the pre-trained cross-tissue imputation models and pre-calculated covariance matrices provided by UTMOST. For the 211 neuroimaging traits in the UKB cohort, we also performed a brain-tissue specific version of UTMOST analysis that only combined brain tissues.

### Comparison with previous GWAS findings

We compared TWAS-significant genes with those identified in the same UKB cohort by MAGMA gene-based association analysis and FUMA functional gene mapping analysis, which can be found in previous GWAS (Supplementary Tables 12 and 15 of Zhao, et al. ^37^ for ROI volumes and Supplementary Tables 14 and 16 of Zhao, et al. ^38^ for DTI parameters, respectively). For each significant gene-trait association, we also explored whether any genetic variant of this gene region (with 1MB window on both sides) had been linked to this neuroimaging trait by checking the smallest p-value in corresponding GWAS. For TWAS-significant genes that were not identified in GWAS, we used NHGRI-EBI GWAS catalog (version 2019-10-14, https://www.ebi.ac.uk/gwas/) to look for their reported associations with brain structure traits and any other traits. We summarized the traits that frequently reported for these genes, such as physical measures (e.g., height, waist-to-hip ratio, heel bone mineral density, body mass index), cognitive functions (such as general cognitive ability, cognitive performance), intelligence, educational attainment, math ability (such as highest math class taken and self-reported math ability), reaction time, neuroticism, neurodegenerative diseases (such as Alzheimer’s disease and Parkinson’s disease), neuropsychiatric disorders (such as major depressive disorder, schizophrenia, and bipolar disorder), coronary artery disease, and mean corpuscular hemoglobin.

### Cross-tissue analysis conditional on the most significant GWAS signal

The TWAS gene expression imputation model can be viewed as a weighted sum of multiple genetic variants. If certain variant has a relatively large weight, the imputed gene expression could be driven by a single GWAS signal. In order to look at how many significant TWAS signals could be dominated by a single genetic variant, we rerun TWAS analysis in UKB cohort conditional on the most significant variant used in the UTMOST imputation model. First, for each reference panel, we considered a simple linear model

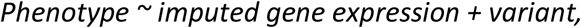

where the variant conditioned on was the most significant variant in previous GWAS of this phenotype in the same UKB cohort. Then, single-tissue conditional p-values of the imputed gene expression were combined by the GBJ test across the 44 GTEx reference panels.

### Enrichment analyses and drug-target lookups

The chromatin interaction enrichments between significant and non-significant genes were tested using the Wilcoxon rank sum test. For the adult neural Promoter Capture Hi-C (PCHi-C), the enrichment of each gene was measured as the number of interactions overlapping gene with CHiCAGO Enrichment Score greater than 5^97^. The enrichment was tested separately in four cell types, including induced pluripotent stem cells (iPSC)-induced excitatory neurons, iPSC-derived hippocampal DG-like neurons, iPSC-induced lower motor neurons, and primary astrocytes. For the high confident interactions of adult and fetal cortex, the enrichment of each gene was measured as the sum of –log_10_(P-value) of all significant interactions overlapping the gene^98^. The drug-target lookups were conducted using the drug-gene associations reported in Wang, et al. ^99^. We focused on nervous system drugs whose Anatomical Therapeutic Chemical code starts with “N” according to the DrugBank database (version 2019-07-02, https://www.drugbank.ca/atc).

### Gene-based TWAS polygenic risk prediction

Gene-based polygenic profiles were created to assess the out-of-sample prediction power of the UKB TWAS results. In this analysis, we used the individual-level phenotype and genetic data, whose processing steps were detailed in previous GWAS^37,38^. The FUSION software and database (http://gusevlab.org/projects/fusion/) were used to impute gene expression levels in UKB, ADNI, HCP, PNC, and PING datasets using individual-level genetic data. We performed imputation for 52 different reference panels (**Supplementary Table 21**). In training data (UKB), we estimated the effect size of each imputed gene expression in a linear regression model, while adjusting for the age (at imaging), age-squared, sex, age-sex interaction, age-squared-sex interaction, as well as the top 40 genetic principle components (PCs) provided by UKB^101^ (Data-Field 22009). For ROI volumes, we also included total brain volume (for ROIs other than total brain volume itself) as a covariate. The gene-based PRS were generated in testing data by summarizing across imputed gene expressions, weighed by their effect sizes estimated from the training data. We tried a series of p-value thresholds for predictor selection: 1, 0.8, 0.5, 0.4, 0.3, 0.2, 0.1, 0.08, 0.05, 0.02, 0.01, 0.001, 1*10^−4^, 1*10^−5^, 1*10^−6^, 1*10^−7^, and 5*10^−8^. Thus, seventeen polygenic profiles were generated for each neuroimaging traits and we reported the best prediction power that can be achieved by a single profile of them in the single reference panel. The association between polygenic profile and trait was estimated and tested in linear regression model, adjusting for the effects of age and sex. The additional phenotypic variation that can be explained by polygenic profile (i.e., the incremental R-squared) was used to measure the prediction power.

### Data availability

The individual-level data used in this work was obtained from five publicly available datasets: the UK Biobank (UKB) study, the Human Connectome Project (HCP) study, the Pediatric Imaging, Neurocognition, and Genetics (PING) study, the Philadelphia Neurodevelopmental Cohort (PNC) study, and the Alzheimer’s Disease Neuroimaging Initiative (ADNI) study. The GWAS summary statistics of UKB study have been shared at https://github.com/BIG-S2/GWAS, and the summary statistics of other validation datasets will also be shared at https://github.com/BIG-S2/GWAS upon acceptance of this paper. We also used the summary-level data of ENIGMA2 and ENIGMA-CHARGE collaboration, which can be obtained at http://enigma.ini.usc.edu/research/. In addition, we used other 11 sets of publicly available GWAS summary statistics shared by several GWAS databases. These data resources were summarized in **Supplementary Table 23**.

### Code availability

We made use of publicly available software and tools, especially the UTMOST (https://github.com/Joker-Jerome/UTMOST) and the FUSION (http://gusevlab.org/projects/fusion/). All codes used to generate results that are reported in this paper are available upon request.

