## Supplementary_information for "Transcriptome-wide association analysis of 211 neuroimaging traits identifies new genes for brain structures and yields insights into the gene-level pleiotropy with other complex traits"

November 11, 2019

#### 1 Supplementary figures

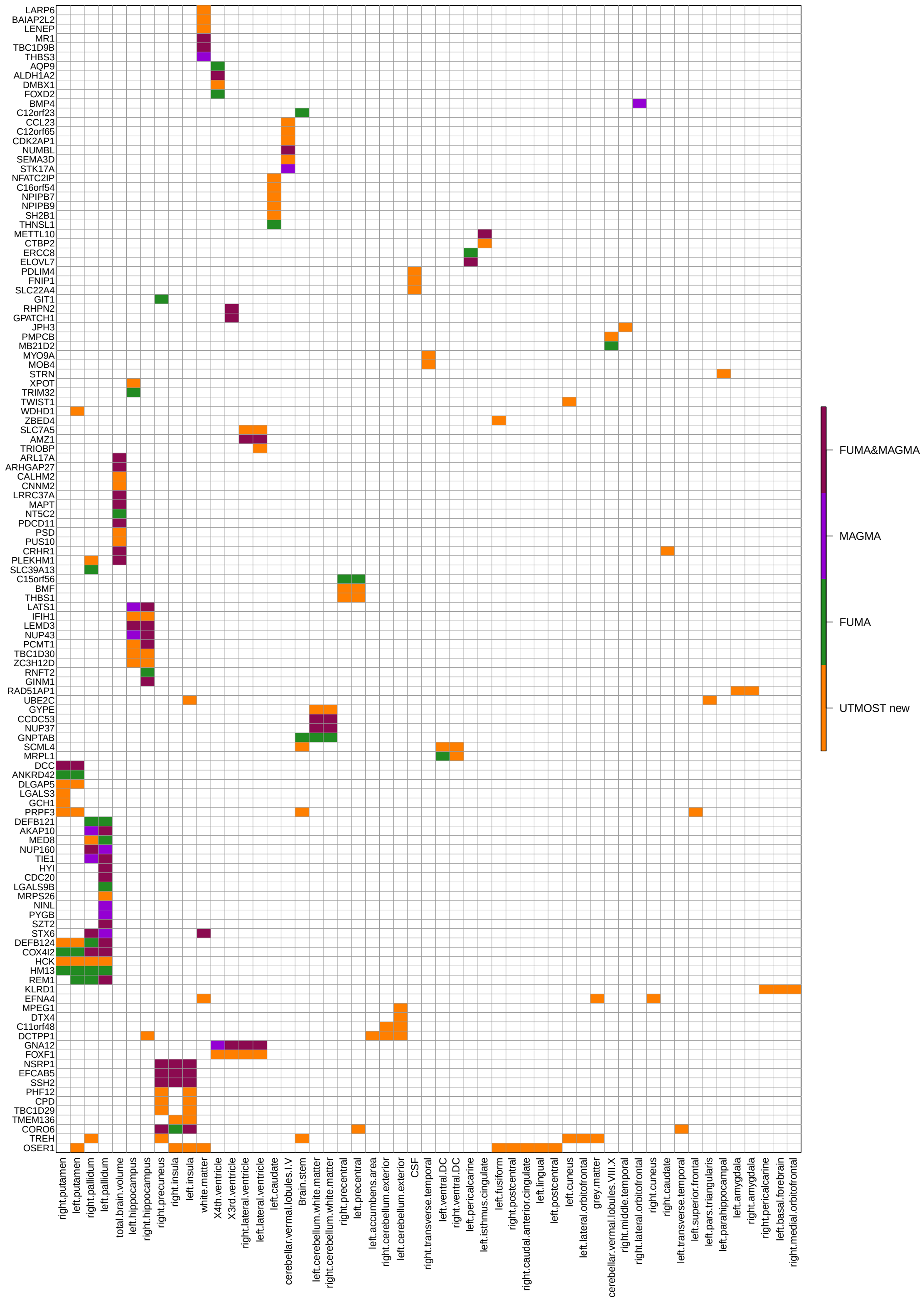


**Supplementary Figure 1:** Significant gene-trait associations discovered in UKB cross-tissue TWAS analysis of ROI volumes (n=19,629 subjects). FUMA: associations identified in FUMA; MAGMA: associations identified in MAGMA; FUMA&MAGMA: associations identified in both FUMA and MAGMA analysis; UTMOST new: novel associations identified in cross-tissue TWAS analysis.

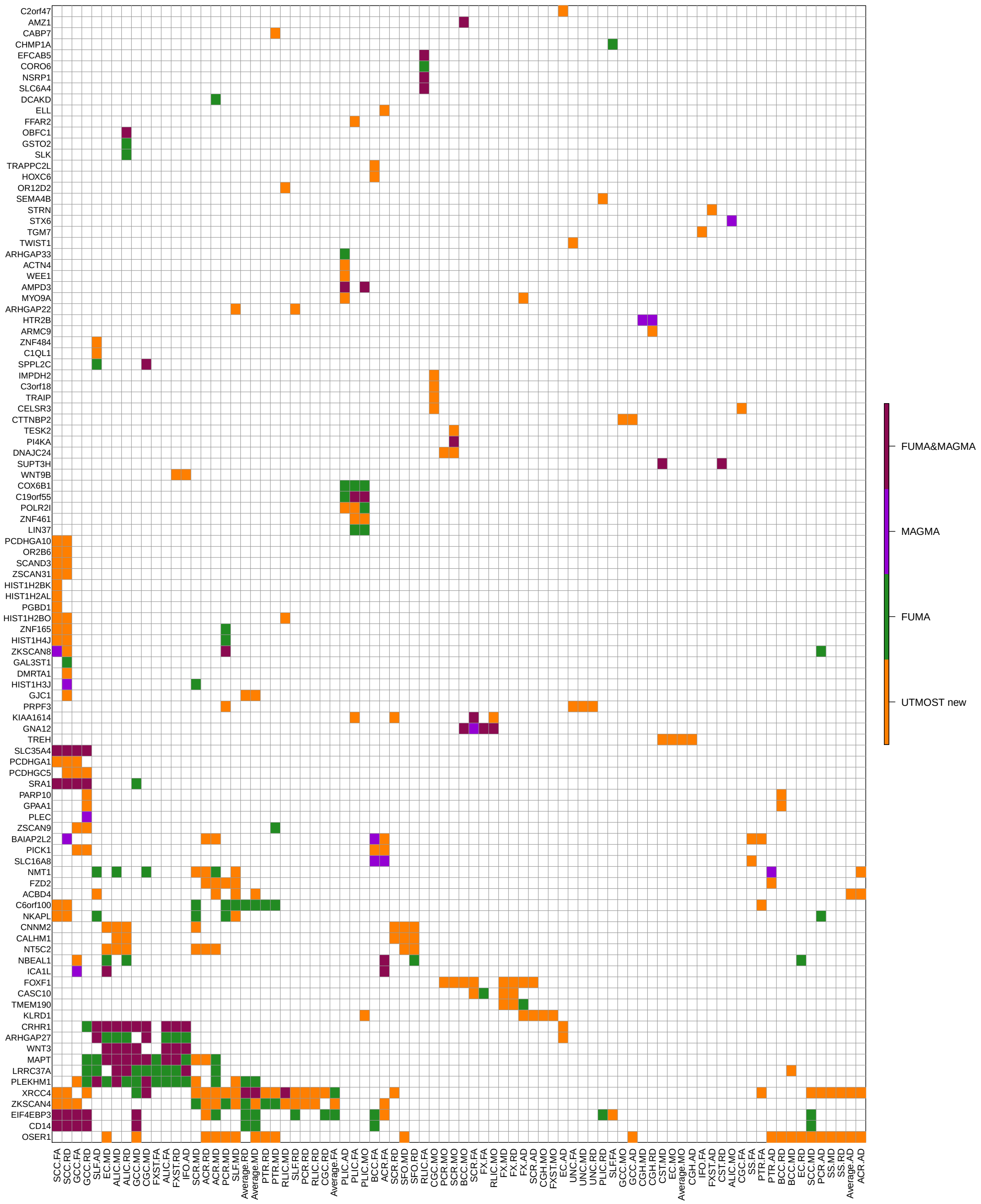

**Supplementary Figure 2:** Significant gene-trait associations discovered in UKB cross-tissue TWAS analysis of DTI parameters (n=17,706 subjects). FUMA: associations identified in FUMA; MAGMA: associations identified in MAGMA; FUMA&MAGMA: associations identified in both FUMA and MAGMA analysis; UTMOST new: novel associations identified in cross-tissue TWAS analysis.

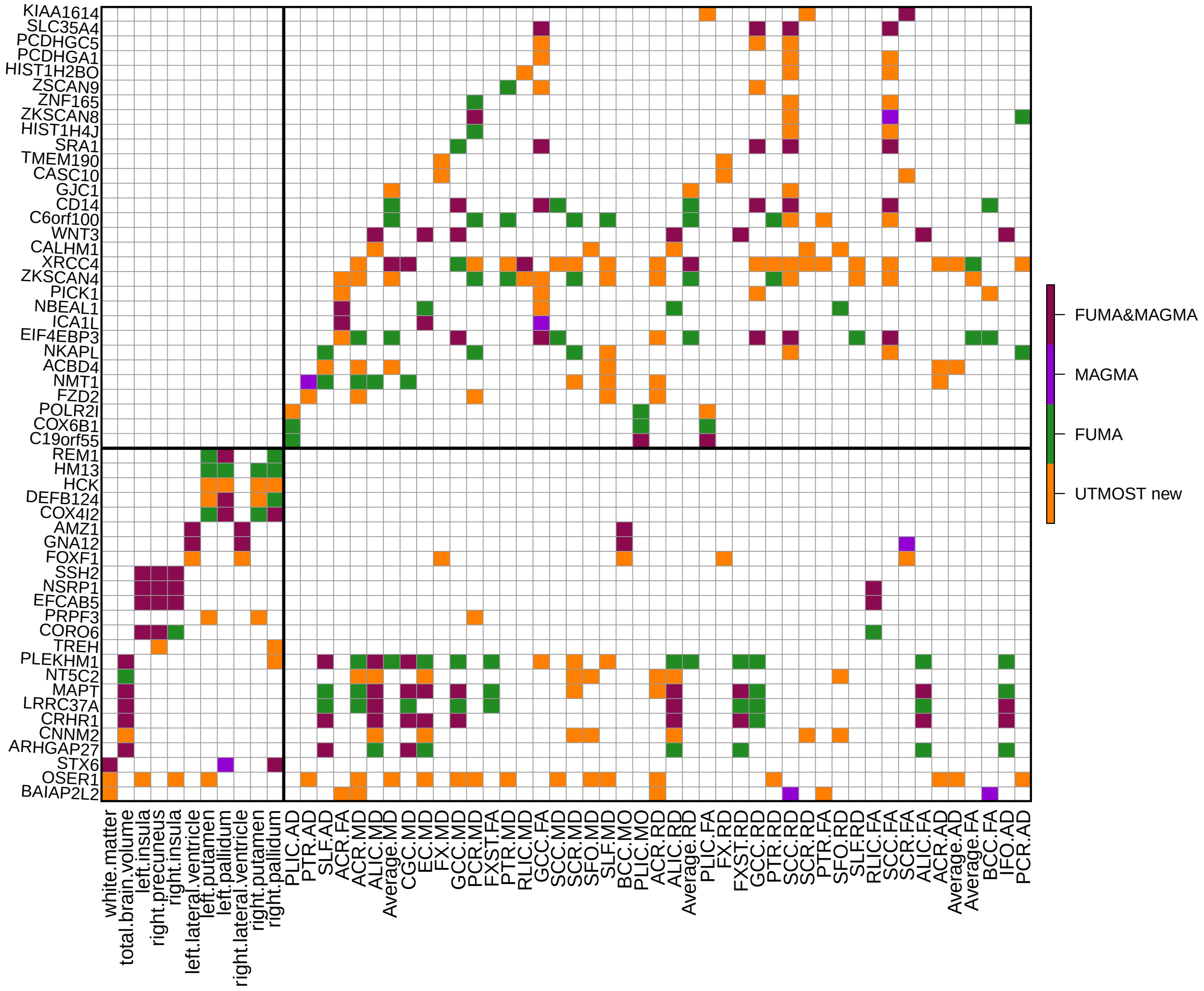

**Supplementary Figure 3:** Selected significant gene-trait associations discovered in UKB cross-tissue TWAS analysis of 211 neuroimaging traits (n=19,629 subjects for ROI volumes and 17,706 for DTI parameters). FUMA: associations identified in FUMA; MAGMA: associations identified in MAGMA; FUMA&MAGMA: associations identified in both FUMA and MAGMA analysis; UTMOST new: novel associations identified in cross-tissue TWAS analysis.

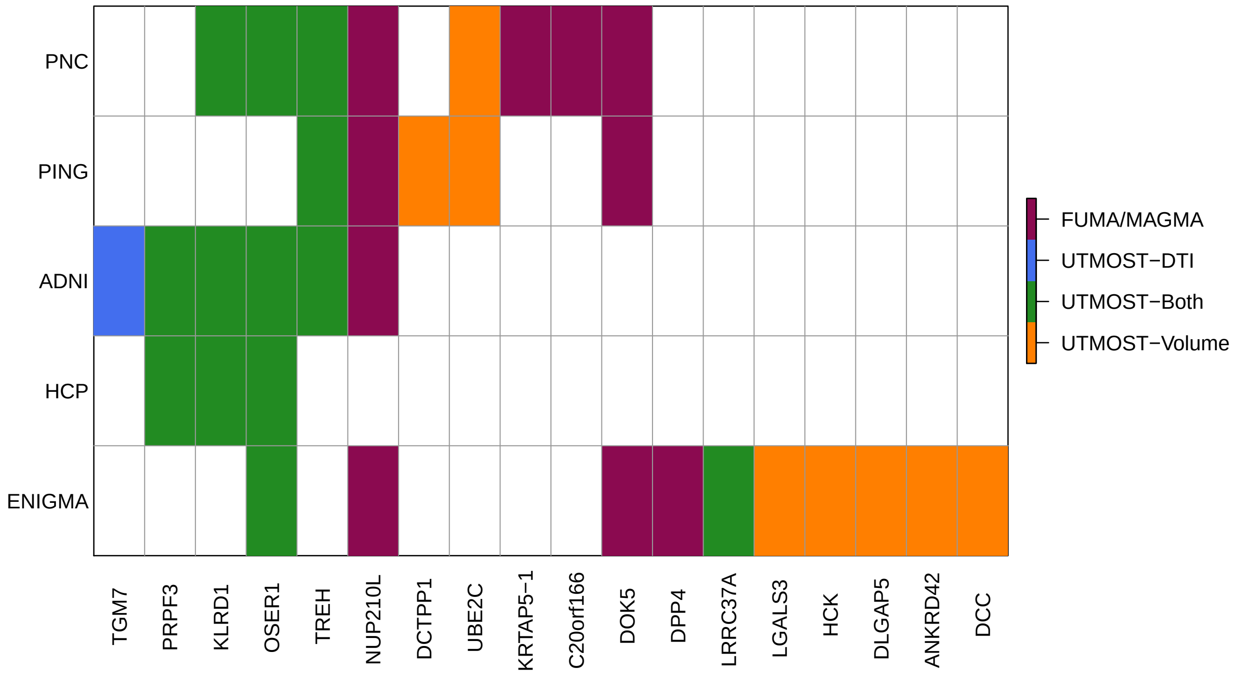


**Supplementary Figure 4:** UKB significant genes that can be validated in one or more of the five validation datasets (n=537 subjects for PNC, 860 subjects for ADNI, 461 subjects for PING, 334 subjects for HCP, and 13,193 subjects for ENIGMA). FUMA/MAGMA: genes identified in FUMA or MAGMA analysis; UTMOST-DTI: genes identified in cross-tissue TWAS analysis for DTI parameters; UTMOST-Volume: genes identified in cross-tissue TWAS analysis for ROI volumes; UTMOST-Both: genes identified in cross-tissue TWAS analysis for both DTI parameters and ROI volumes.

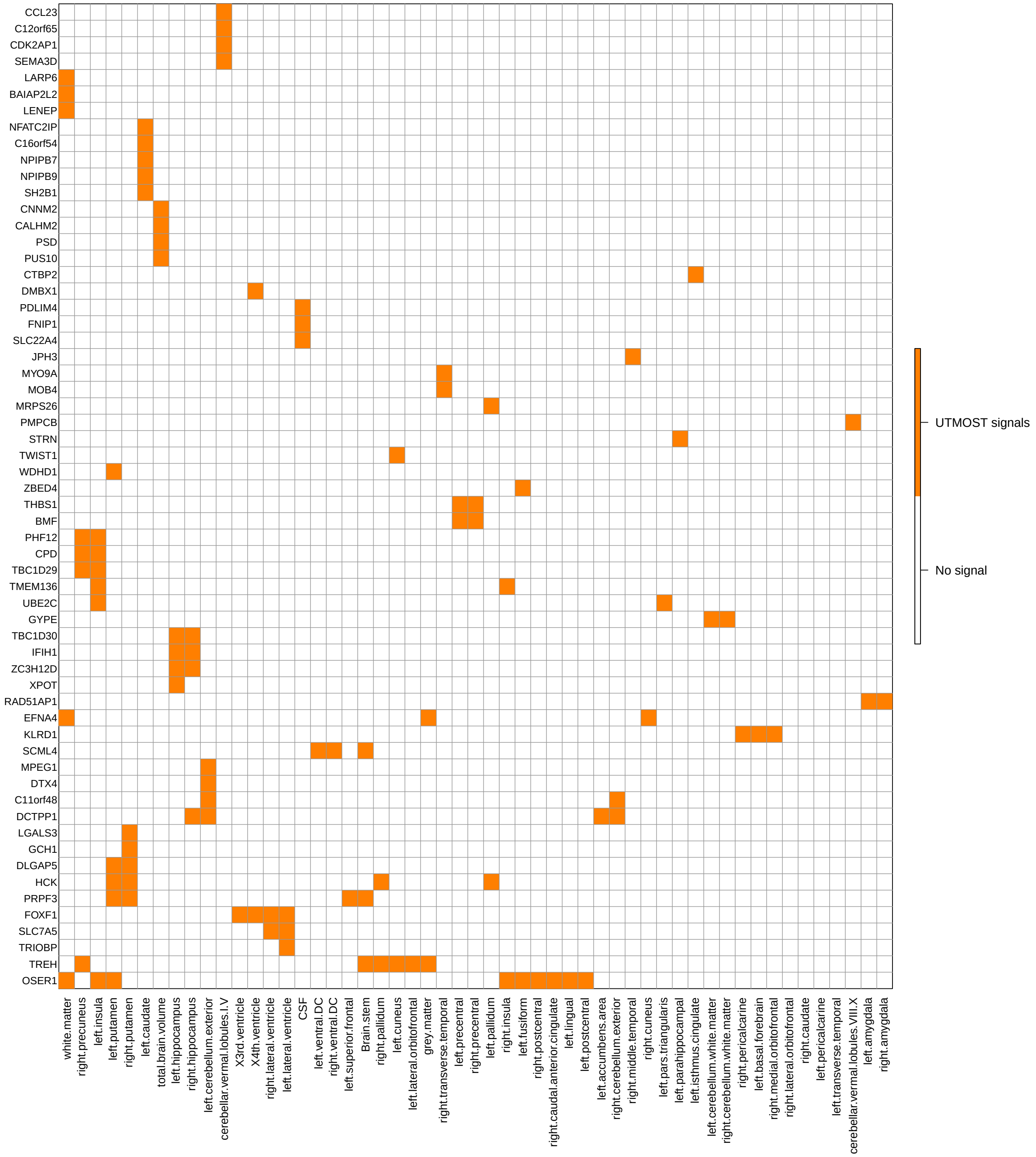


**Supplementary Figure 5:** Significant genes and their associations identified in UKB cross-tissue TWAS analysis of ROI volumes (n=19,629 subjects) but not in previous GWAS of the same dataset.

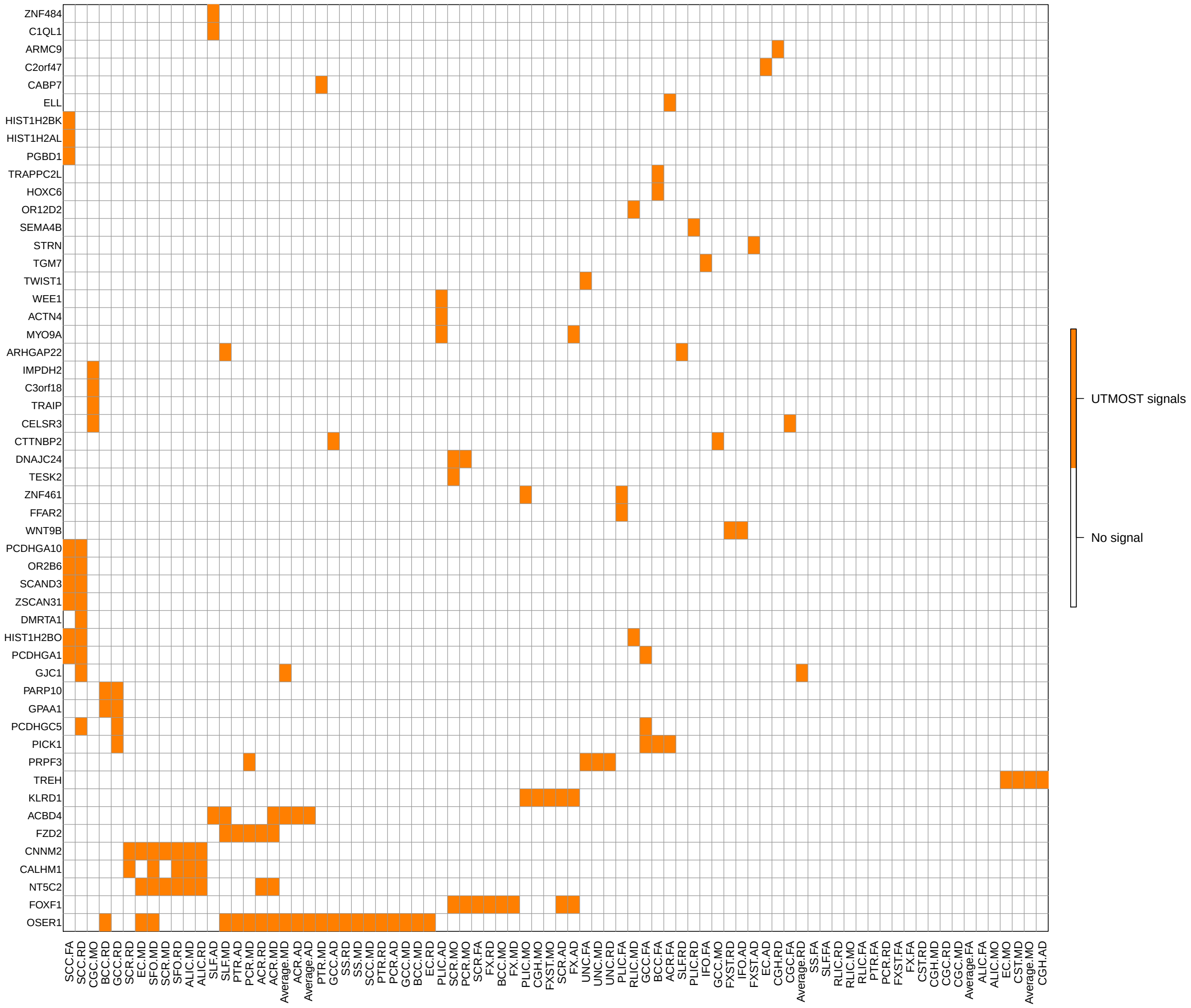


**Supplementary Figure 6:** Significant genes and their associations identified in UKB cross-tissue TWAS analysis of ROI volumes (n=17,706 subjects) but not in previous GWAS of the same dataset.

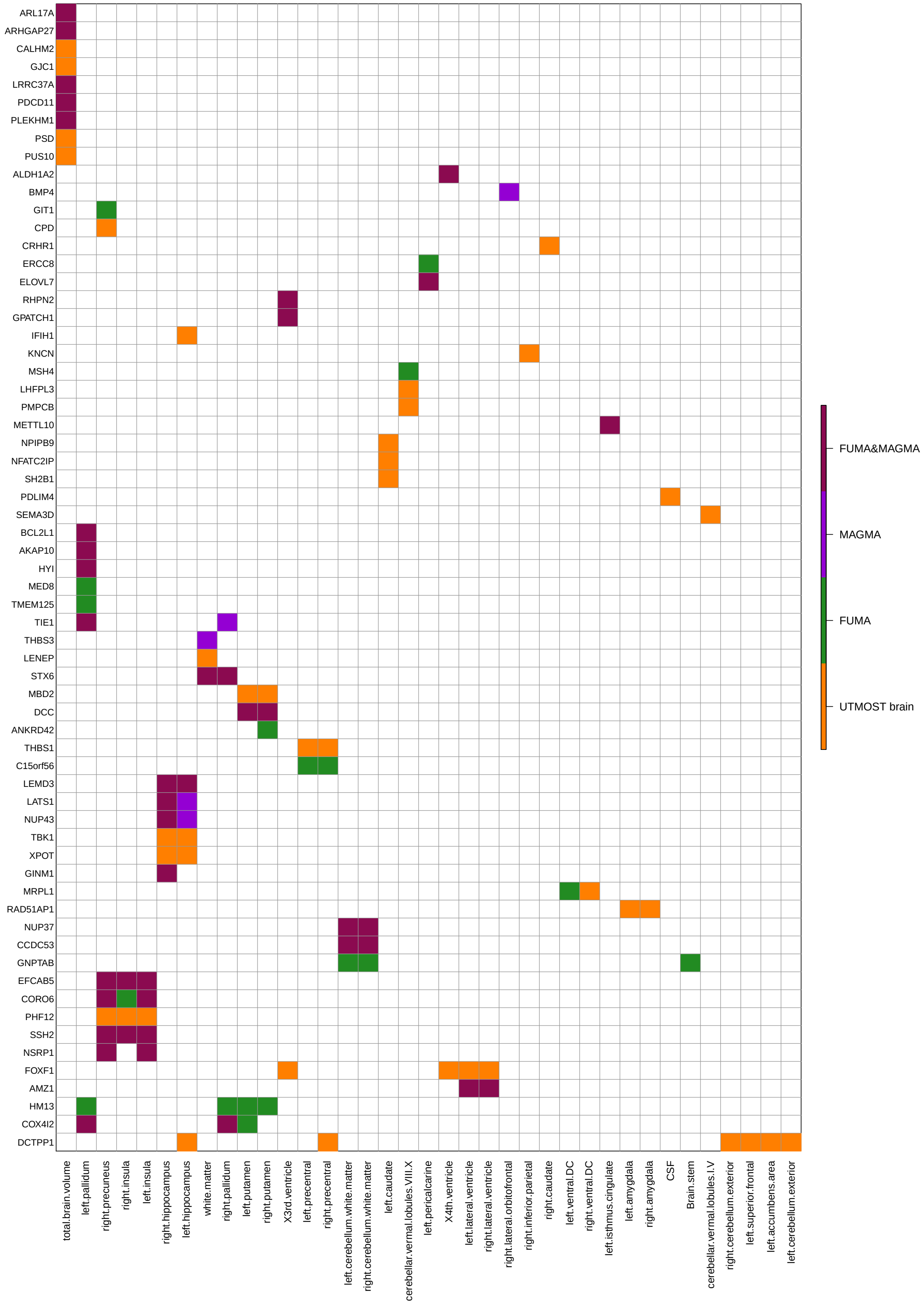


**Supplementary Figure 7:** Significant gene-trait associations discovered in UKB brain tissue-specific TWAS analysis of ROI volumes (n=19,629 subjects). FUMA: associations identified in FUMA; MAGMA: associations identified in MAGMA; FUMA&MAGMA: associations identified in both FUMA and MAGMA analysis; UTMOST brain: novel associations identified in brain tissue-specific TWAS analysis.

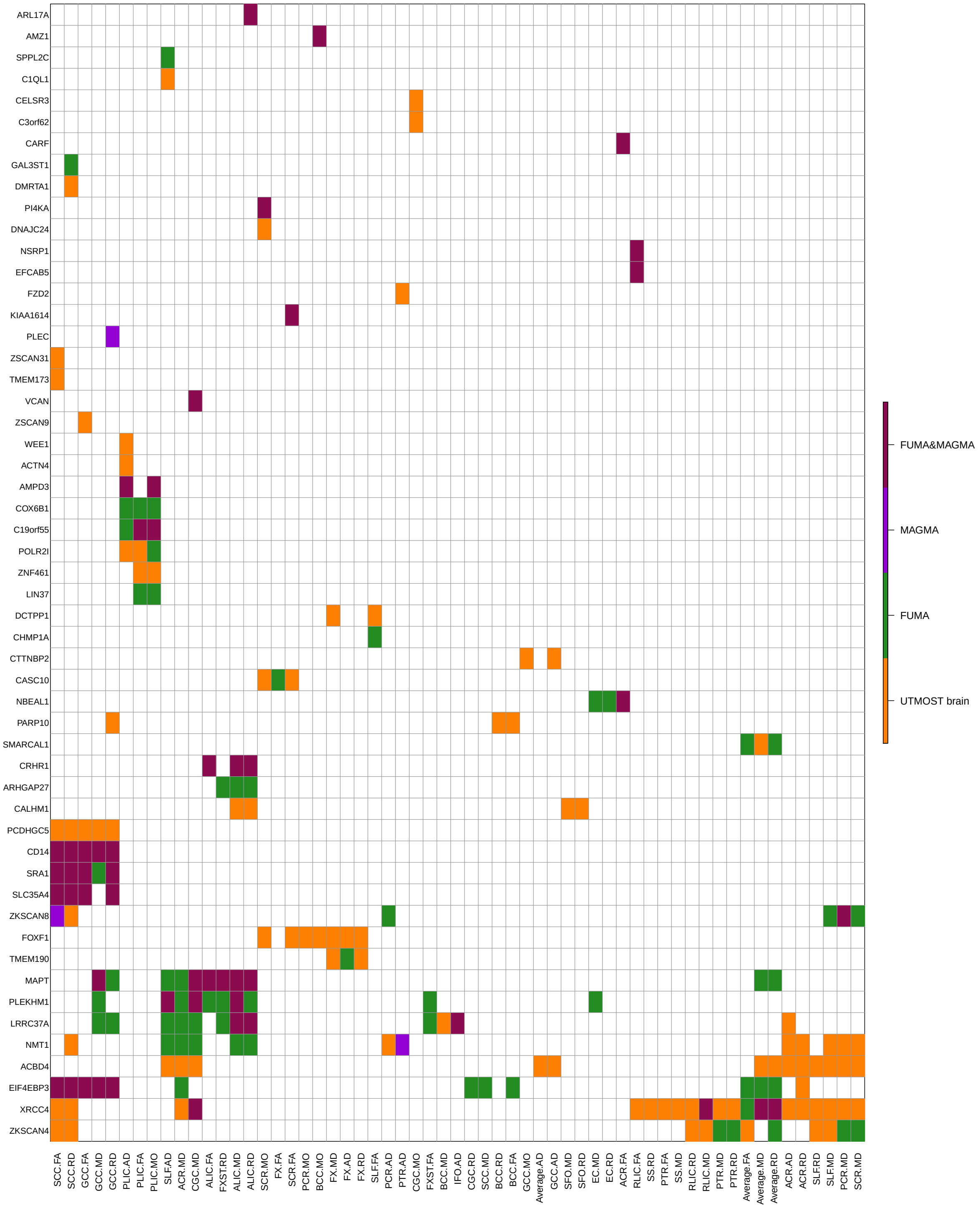


**Supplementary Figure 8:** Significant gene-trait associations discovered in UKB brain tissue-specific TWAS analysis of DTI parameters (n=17,706 subjects). FUMA: associations identified in FUMA; MAGMA: associations identified in MAGMA; FUMA&MAGMA: associations identified in both FUMA and MAGMA analysis; UTMOST brain: novel associations identified in brain tissue-specific TWAS analysis.

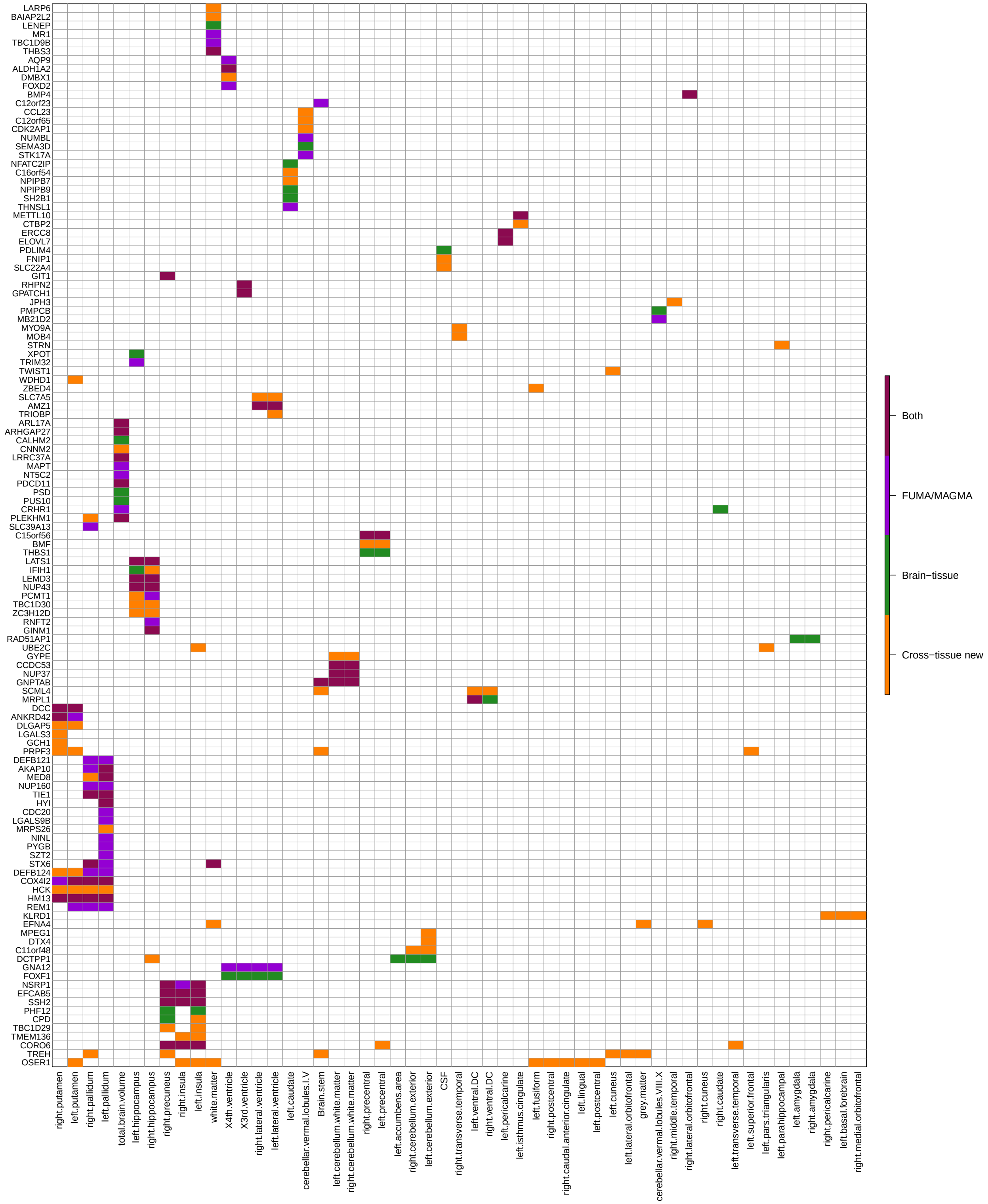


**Supplementary Figure 9:** Significant gene-trait associations discovered in UKB cross-tissue TWAS analysis of ROI volumes (n=19,629 subjects). Brain-tissue: associations identified in UKB brain tissue-specific analysis; Both: associations identified in both UKB brain tissue-specific analysis and FUMA or MAGMA analysis; Cross-tissue new: novel associations identifed in cross-tissue TWAS analysis.

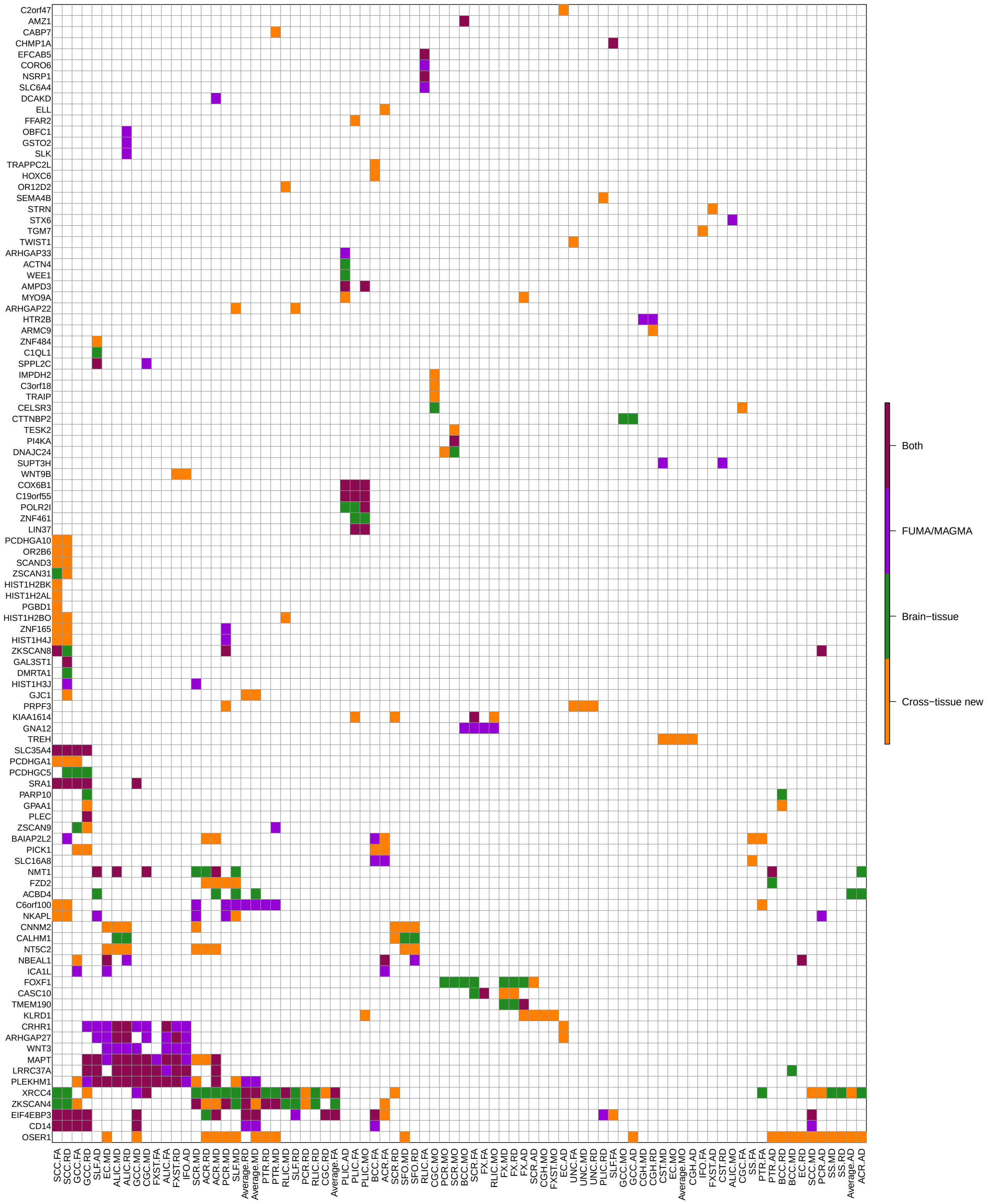


**Supplementary Figure 10:** Significant gene-trait associations discovered in UKB cross-tissue TWAS analysis of DTI parameters (n=17,706 subjects). Brain-tissue: associations identified in UKB brain tissue-specific analysis; Both: associations identified in both UKB brain tissue-specific analysis and FUMA or MAGMA analysis; Cross-tissue new: novel associations identifed in cross-tissue TWAS analysis.

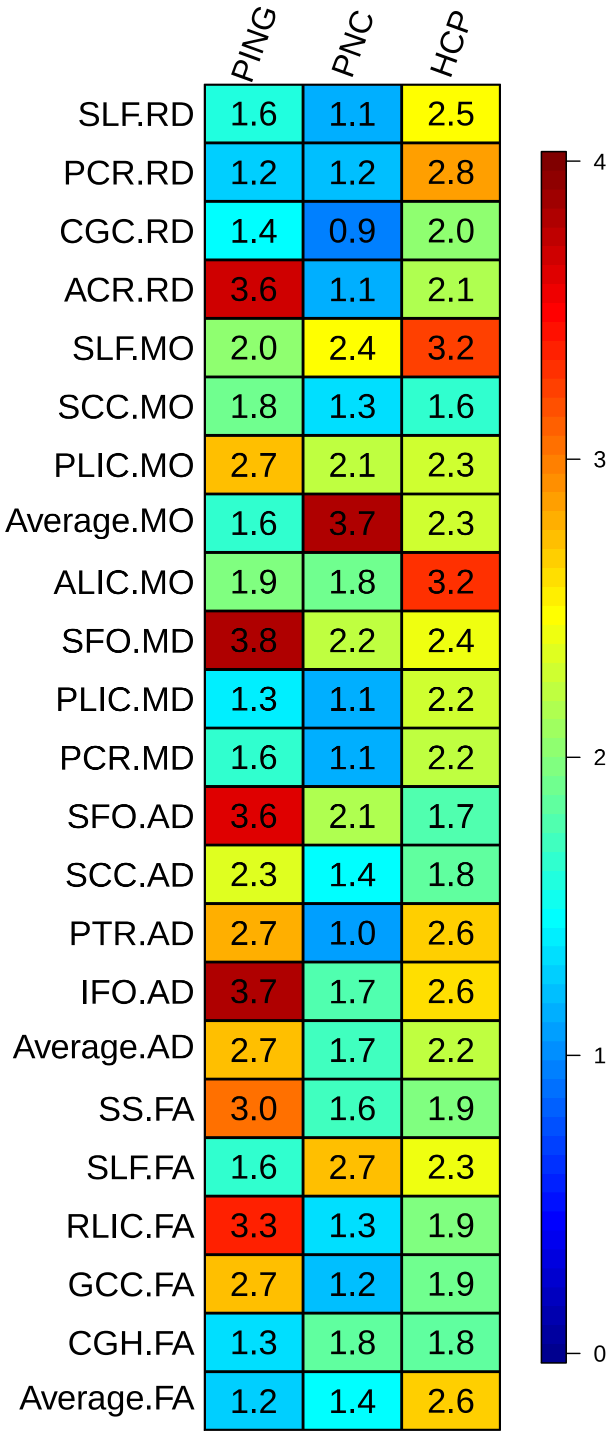


**Supplementary Figure 11:** Prediction accuracy (incremental R-squared) of significant gene-based polygenic risk scores constructed by UKB-derived TWAS summary statistics (n=17,706 subjects) on the three independent datasets (PING, PNC, HCP). We display the 23 DTI parameters that are significant in all the three datasets after the Bonferroni correction.

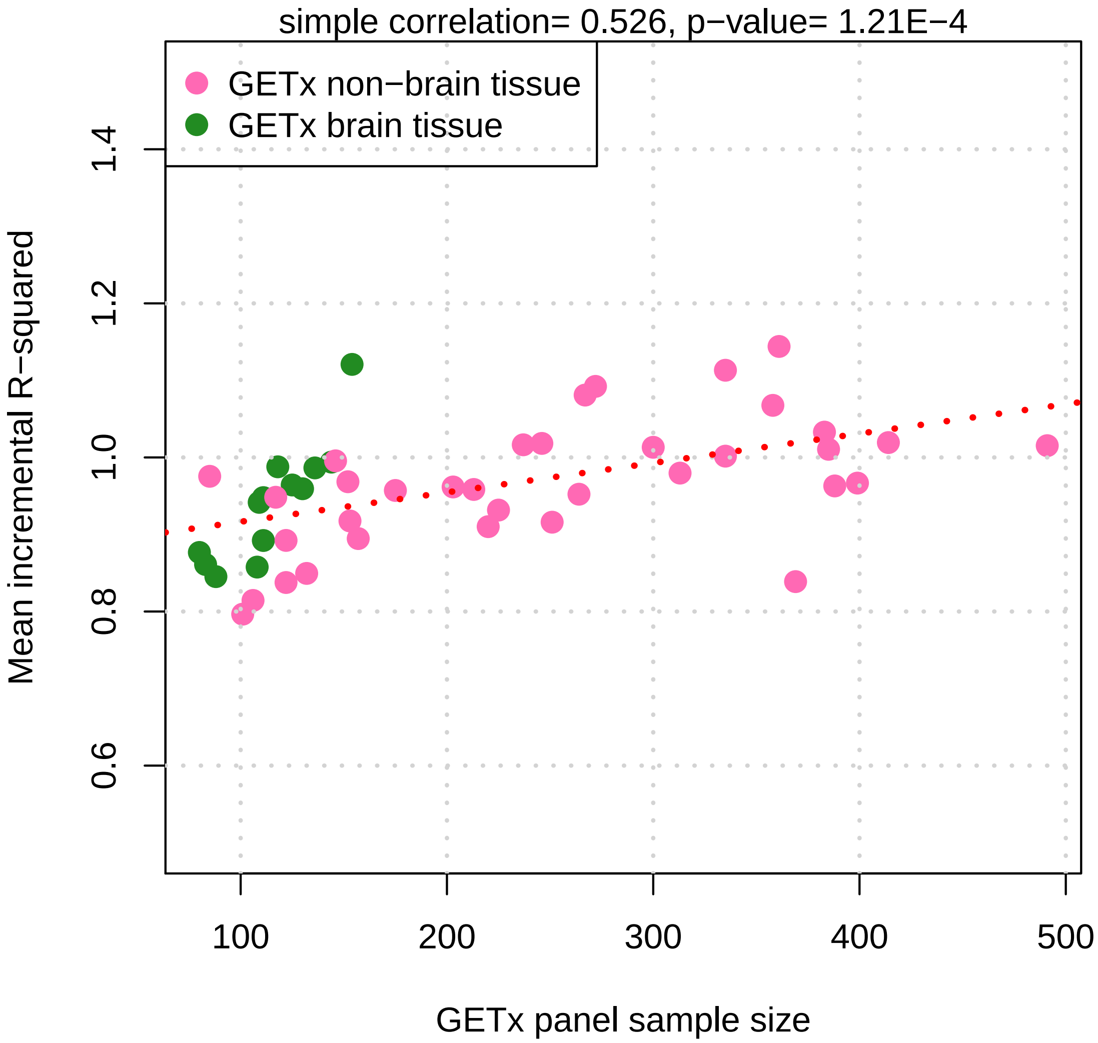


**Supplementary Figure 12:** Relationship between mean incremental prediction R-squared of significant gene-based polygenic risk scores constructed on each GTEx reference panel by UKB-derived GWAS summary statistics (n=19,629 subjects for ROI volumes and 17,706 for DTI parameters, respectively) and the sample size of reference panels.

### 2 Supplementary tables

All supplementary tables can be found in the zip file, here is a list the header of each table.

**Supplementary Table 1:** List of significant gene-trait associations discovered in UKB cross-tissue TWAS analysis of 211 neuroimaging traits (n=19,629 subjects for ROI volumes and 17,706 for DTI parameters). The p-values are raw p-values of Generalized Berk-Jones (GBJ) test (`https://cran.r-project.org/web/packages/GBJ/vignettes/GBJ_tutorial.html`) generated by UTMOST software (`https://github.com/Joker-Jerome/UTMOST/`). We refer $1.37 \times 10^{-8}$ (that is, 0.05/17,290/211, adjusted for all candidate genes and traits performed) as the significance threshold.

**Supplementary Table 2:** List of significant gene-trait associations identified in ADNI cross-tissue TWAS analysis of 101 ROI volumes (n=860 subjects). The p-values are raw p-values of Generalized Berk-Jones (GBJ) test (`https://cran.r-project.org/web/packages/GBJ/vignettes/GBJ_tutorial.html`) generated by UTMOST software (`https://github.com/Joker-Jerome/UTMOST/`). We refer $2.86 \times 10^{-8}$ (that is, 0.05/17,290/101, adjusted for all candidate genes and traits performed) as the significance threshold.

**Supplementary Table 3:** List of significant gene-trait associations identified in ENIGMA cross-tissue TWAS analysis of 8 ROI volumes (n=13,193 subjects). The p-values are raw p-values of Generalized Berk-Jones (GBJ) test (`https://cran.r-project.org/web/packages/GBJ/vignettes/GBJ_tutorial.html`) generated by UTMOST software (`https://github.com/Joker-Jerome/UTMOST/`). We refer $3.61 \times 10^{-7}$ (that is, 0.05/17,290/8, adjusted for all candidate genes and traits performed) as the significance threshold.

**Supplementary Table 4:** List of significant gene-trait associations discovered in HCP cross-tissue TWAS analysis of 211 neuroimaging traits (n=334 subjects for ROI volumes and 319 for DTI parameters). The p-values are raw p-values of Generalized Berk-Jones (GBJ) test (`https://cran.r-project.org/web/packages/GBJ/vignettes/GBJ_tutorial.html`) generated by UTMOST software (`https://github.com/Joker-Jerome/UTMOST/`). We refer $1.37 \times 10^{-8}$ (that is, 0.05/17,290/211, adjusted for all candidate genes and traits performed) as the significance threshold.

**Supplementary Table 5:** List of significant gene-trait associations discovered in PING cross-tissue TWAS analysis of 211 neuroimaging traits (n=461 subjects for ROI volumes and 444 for DTI parameters). The p-values are raw p-values of Generalized Berk-Jones (GBJ) test (`https://cran.r-project.org/web/packages/GBJ/vignettes/GBJ_tutorial.html`) generated by UTMOST software (`https://github.com/Joker-Jerome/UTMOST/`). We refer $1.37 \times 10^{-8}$ (that is, 0.05/17,290/211, adjusted for all candidate genes and traits performed) as the significance threshold.

**Supplementary Table 6:** List of significant gene-trait associations discovered in PNC cross-tissue TWAS analysis of 211 neuroimaging traits (n=537 subjects for ROI volumes and 520 for DTI parameters). The p-values are raw p-values of Generalized Berk-Jones (GBJ) test (`https://cran.r-project.org/web/packages/GBJ/vignettes/GBJ_tutorial.html`) generated by UTMOST software (`https://github.com/Joker-Jerome/UTMOST/`). We refer $1.37 \times 10^{-8}$ (that is, 0.05/17,290/211, adjusted for all candidate genes and traits performed) as the significance threshold.

**Supplementary Table 7:** List of significant genes identified in UKB cross-tissue TWAS analysis of 211 neuroimaging traits (n=19,629 subjects for ROI volumes and 17,706 for DTI parameters) that were not discovered in previous GWAS of the same dataset.

**Supplementary Table 8:** Associated traits of the 90 TWAS-significant genes (listed in Supplementary Table 7) that have previously been reported in the NHGRI-EBI GWAS catalog (version 2019-10-14, www.ebi.ac.uk/gwas/).

**Supplementary Table 9:** List of significant gene-trait associations discovered in UKB brain tissue-specific TWAS analysis of 211 neuroimaging traits (n=19,629 subjects for ROI volumes and 17,706 for DTI parameters). The p-values are raw p-values of Generalized Berk-Jones (GBJ) test (`https://cran.r-project.org/web/packages/GBJ/vignettes/GBJ_tutorial.html`) generated by UTMOST software (`https://github.com/Joker-Jerome/UTMOST/`). We refer $1.37 \times 10^{-8}$ (that is, 0.05/17,290/211, adjusted for all candidate genes and traits performed) as the significance threshold.

**Supplementary Table 10:** The most significant GWAS variant-level signal corresponding to the gene-trait associations discovered in UKB cross-tissue TWAS analysis of 211 neuroimaging traits (n=19,629 subjects for ROI volumes and 17,706 for DTI parameters)

**Supplementary Table 11:** The most significant GWAS variant-level signal corresponding to the gene-trait associations discovered in UKB brain tissue-specific TWAS analysis of 211 neuroimaging traits (n=19,629 subjects for ROI volumes and 17,706 for DTI parameters)

**Supplementary Table 12:** Conditional analysis of significant gene-trait associations discovered in UKB cross-tissue TWAS analysis of 211 neuroimaging traits (n=19,629 subjects for ROI volumes and 17,706 for DTI parameters). The p-values are raw conditional p-values of Generalized Berk-Jones (GBJ) test (`https://cran.r-project.org/web/packages/GBJ/vignettes/GBJ_tutorial.html`) generated by UTMOST software (`https://github.com/Joker-Jerome/UTMOST/`). We refer $1.37 \times 10^{-8}$ (that is, 0.05/17,290/211, adjusted for all candidate genes and traits performed) as the significance threshold.

**Supplementary Table 13:** Conditional analysis of significant gene-trait associations discovered in UKB brain tissue-specific TWAS analysis of 211 neuroimaging traits (n=19,629 subjects for ROI volumes and 17,706 for DTI parameters). The p-values are raw conditional p-values of Generalized Berk-Jones (GBJ) test (`https://cran.r-project.org/web/packages/GBJ/vignettes/GBJ_tutorial.html`) generated by UTMOST software (`https://github.com/Joker-Jerome/UTMOST/`). We refer $1.37 \times 10^{-8}$ (that is, 0.05/17,290/211, adjusted for all candidate genes and traits performed) as the significance threshold.

**Supplementary Table 14:** List of significant gene-trait associations discovered in UKB cross-tissue TWAS analysis of 11 cognitive and mental health traits. (sample sizes and data resources can be found in Supplementary Table 23). The p-values are raw p-values of Generalized Berk-Jones (GBJ) test (`https://cran.r-project.org/web/packages/GBJ/vignettes/GBJ_tutorial.html`) generated by UTMOST software (`https://github.com/Joker-Jerome/UTMOST/`). We refer $2.63 \times 10^{-7}$ (that is, 0.05/17,290/11, adjusted for all candidate genes and traits performed) as the significance threshold.

**Supplementary Table 15:** Enrichment analysis of genes identified for 211 neuroimaging traits (n=19,629 subjects for ROI volumes and 17,706 for DTI parameters) and 11 cognitive and mental health traits. (sample sizes and data resources can be found in Supplementary Table 23). The p-values are raw p-values of Wilcoxon rank test.

**Supplementary Table 16:** Enrichment analysis of genes identified for 211 neuroimaging traits (n=19,629 subjects for ROI volumes and 17,706 for DTI parameters) and 11 cognitive and mental health traits. (sample sizes and data resources can be found in Supplementary Table 23). The p-values are adjusted p-values after correcting for multiple testing by Benjamini-Hochberg (B-H) procedure at 0.05 level.

**Supplementary Table 17:** Drug-target lookups of genes identified for 11 cognitive and mental health traits (sample sizes and data resources can be found in Supplementary Table 23).

**Supplementary Table 18:** Drug-target lookups of genes identified for 211 neuroimaging traits (n=19,629 subjects for ROI volumes and 17,706 for DTI parameters).

**Supplementary Table 19:** Prediction accuracy (incremental R-squared) and p-value of gene-based polygenic risk scores constructed by UKB-derived TWAS summary statistics (n=19,629 subjects) on the four independent datasets. The p-values are asymptotic p-values of two-sided t-test statistics in linear regression.

**Supplementary Table 20:** Prediction accuracy (incremental R-squared) and p-value of gene-based polygenic risk scores constructed by UKB-derived GWAS summary statistics (n=17,706 subjects) on the three independent datasets. The p-values are asymptotic p-values of two-sided t-test statistics in linear regression.

**Supplementary Table 21:** Mean prediction accuracy (incremental R-squared) of gene-based polygenic risk scores constructed using each reference panel (n=19,629 subjects for ROI volumes and 17,706 for DTI parameters).

**Supplementary Table 22:** Sample size of each neuroimaging GWAS and IDs of neuroimaging traits.

**Supplementary Table 23:** Sources of the 11 sets of publicly available GWAS summary statistics of cognitive and mental health traits used in this study.

### 3 Supplementary Note

#### PING Methods

Part of the data used in the preparation of this article were obtained from the Pediatric Imaging, Neurocognition and Genetics (PING) Study database (http://ping.chd.ucsd.edu/). PING was launched in 2009 by the National Institute on Drug Abuse (NIDA) and the Eunice Kennedy Shriver National Institute Of Child Health & Human Development (NICHD) as a 2-year project of the American Recovery and Reinvestment Act. The primary goal of PING has been to create a data resource of highly standardized and carefully curated magnetic resonance imaging (MRI) data, comprehensive genotyping data, and developmental and neuropsychological assessments for a large cohort of developing children aged 3 to 20 years. The scientific aim of the project is, by openly sharing these data, to amplify the power and productivity of investigations of healthy and disordered development in children, and to increase understanding of the origins of variation in neurobehavioral phenotypes. For up-to-date information, see http://ping.chd.ucsd.edu/.

#### ADNI Methods

Data used in the preparation of this article were obtained from the Alzheimer's Disease Neuroimaging Initiative (ADNI) database (http://adni.loni.usc.edu). The ADNI was launched in 2003 by the National Institute on Aging (NIA), the National Institute of Biomedical Imaging and Bioengineering (NIBIB), the Food and Drug Administration (FDA), private pharmaceutical companies and non-profit organizations, as a 60 million, 5-year public-private partnership. The primary goal of ADNI has been to test whether serial magnetic resonance imaging (MRI), positron emission tomography (PET), other biological markers, and clinical and neuropsychological assessment can be combined to measure the progression of mild cognitive impairment (MCI) and early Alzheimer's disease (AD). Determination of sensitive and specific markers of very early AD progression is intended to aid researchers and clinicians to develop new treatments and monitor their effectiveness, as well as lessen the time and cost of clinical trials.

The Principal Investigator of this initiative is Michael W. Weiner, MD, VA Medical Center and University of California – San Francisco. ADNI is the result of efforts of many co-investigators from a broad range of academic institutions and private corporations, and subjects have been recruited from over 50 sites across the U.S. and Canada. The initial goal of ADNI was to recruit 800 subjects but ADNI has been followed by ADNI-GO and ADNI-2. To date these three protocols have recruited over 1500 adults, ages 55 to 90, to participate in the research, consisting of cognitively normal older individuals, people with early or late MCI, and people with early AD. The follow up duration of each group is specified in the protocols for ADNI-1, ADNI-2 and ADNI-GO. Subjects originally recruited for ADNI-1 and ADNI-GO had the option to be followed in ADNI-2. For up-to-date information, see www.adni-info.org.

#### Pediatric Imaging, Neurocognition and Genetics (PING) Authors

Connor McCabe[1], Linda Chang[2], Natacha Akshoomoff[3], Erik Newman[1], Thomas Ernst[2], Peter Van Zijl[4], Joshua Kuperman[5], Sarah Murray[6], Cinnamon Bloss[6], Mark Appelbaum[1], Anthony Gamst[1], Wesley Thompson[3], Hauke Bartsch[5].

#### Alzheimer's Disease Neuroimaging Initiative (ADNI) Authors

Michael Weiner[7], Paul Aisen[1], Ronald Petersen[8], Clifford R. Jack Jr[8], William Jagust[9], John Q. Trojanowki[10], Arthur W. Toga[11], Laurel Beckett[12], Robert C. Green[13], Andrew J. Saykin[14], John Morris[15], Leslie M. Shaw[10], Zaven Khachaturian[16], Greg Sorensen[17], Maria Carrillo[18], Lew Kuller[19], Marc Raichle[15], Steven Paul[20], Peter Davies[21], Howard Fillit[22], Franz Hefti[23], Davie Holtzman[15], M. Marcel Mesulman[24], William Potter[25], Peter J. Snyder[26], Adam Schwartz[27], Tom Montine[28], Ronald G. Thomas[1], Michael Donohue[1], Sarah Walter[1], Devon Gessert[1], Tamie Sather[1], Gus Jiminez[1], Danielle Harvey[12], Matthew Bernstein[8], Nick Fox[29], Paul Thompson[11], Norbert Schuff[7], Charles DeCarli[12], Bret Borowski[8], Jeff Gunter[8], Matt Senjem[8], Prashanthi Vemuri[8], David Jones[8], Kejal Kantarci[8], Chad Ward[8], Robert A. Koeppe[30], Norm Foster[31], Eric M. Reiman[32], Kewei Chen[32], Chet Mathis[19], Susan Landau[9], Nigel J. Cairns[15], Erin Householder[15], Lisa Taylor-Reinwald[15], Virginia M.Y. Lee[10], Magdalena Korecka[10], Michal Figurski[10], Karen Crawford[11], Scott Neu[11], Tatiana M. Foroud[14], Steven Potkin[33], Li Shen[14], Kelley Faber[14], Sungeun Kim[14], Kwangsik Nho[14], Leon Thal[1], Richard Frank[34], Neil Buckholtz[35], Marilyn Albert[36], John Hsiao[35].

[1]UC San Diego, La Jolla, CA 92093, USA. [2]U Hawaii, Honolulu, HI 96822, USA. [3]Department of Psychiatry, University of California, San Diego, La Jolla, California 92093, USA. [4]Kennedy Krieger Institute, Baltimore, MD 21205, USA. [5]Multimodal Imaging Laboratory, Department of Radiology, University of California San Diego, La Jolla, California 92037, USA. [6]Scripps Translational Science Institute, La Jolla, CA 92037, USA. [7]UC San Francisco, San Francisco, CA 94143, USA. [8]Mayo Clinic, Rochester, MN 55905, USA. [9]UC Berkeley, Berkeley, CA 94720-5800, USA. [10]U Pennsylvania, Philadelphia, PA 19104, USA. [11]USC, University of Southern California, Los Angeles, CA 90033, USA. [12]UC Davis, Davis, CA 95616, USA. [13]Brigham and Women s Hospital/Harvard Medical School, Boston MA 02115, USA. [14]Indiana University, Indianapolis, IN 46202-5143, USA. [15]Washington

University St. Louis, St. Louis, MO 63130, USA. [16]Prevent Alzheimer's Disease 2020, Rockville, MD 20850, USA. [17]Siemens [18]Alzheimer's Association, Chicago, IL 60601, USA. [19]University of Pittsburgh, Pittsburgh, PA 15260, USA. [20]Cornell University, Ithaca, NY 14850, USA. [21]Albert Einstein College of Medicine of Yeshiva University, Bronx, NY 10461, USA. [22]AD Drug Discovery Foundation, New York, NY 10019, USA. [23]Acumen Pharmaceuticals, Livermore, California 94551, USA. [24]Northwestern University, Evanston, IL 60208, USA. [25]National Institute of Mental Health, Bethesda, MD 20892-9663, USA. [26]Brown University, Providence, RI 02912, USA. [27]Eli Lilly, Indianapolis, Indiana 46285, USA. [28]University of Washington, Seattle, WA 98195, USA. [29]University of London, London WC1E 7HU, UK. [30]University of Michigan, Ann Arbor, MI 48109, USA. [31]University of Utah, Salt Lake City, UT 84112, USA. [32]Banner Alzheimer's Institute, Phoenix, AZ 85006, USA. [33]UC Irvine, Irvine, CA 92697, USA. [34]General Electric [35]National Institute on Aging/National Institutes of Health, Bethesda, MD 20892, USA. [36]The Johns Hopkins University, Baltimore, MD 21218, USA.
